# M4-mediated cholinergic transmission is reduced in parkinsonian mice and its restoration alleviates motor deficits and levodopa-induced dyskinesia

**DOI:** 10.1101/2023.08.23.554510

**Authors:** Beatriz E. Nielsen, Christopher P. Ford

## Abstract

The dynamic equilibrium between dopamine (DA) and acetylcholine (ACh) in the dorsal striatum is thought to be essential for motor function, as imbalances in their levels are associated with Parkinson’s disease (PD) and levodopa-induced dyskinesia (LID). While loss of DA leads to enhanced striatal ACh, whether this translates to specific alterations in transmission remains unclear. To address this, we examined how the strength of ACh release and signaling onto direct-pathway medium spiny neurons is altered in parkinsonian mice. Rather than the predicted cholinergic enhancement, we found that the strength of muscarinic M4-receptor mediated transmission was reduced following DA loss, resulting from downregulated receptors and downstream signaling. Despite M4-receptors being thought to mediate anti-kinetic effects, restoring M4-receptor function partially rescued parkinsonian balance and coordination deficits and limited the development of levodopa-induced dyskinetic behaviors, indicating that decreased M4-function contributed to circuit and motor dysfunctions in response to DA loss.

## INTRODUCTION

Parkinson’s disease (PD) is a neurodegenerative movement disorder characterized by the progressive loss of dopamine (DA) neurons of the substantia nigra pars compacta (SNc), resulting in dopaminergic denervation of the dorsal striatum (DSt). The loss of DA drives motor impairments as a result of alterations in striatal circuits, that ultimately lead to imbalances in output from the direct and indirect pathways, comprised of medium spiny neurons expressing either D1-receptors (dMSNs) or D2-receptors (iMSNs) (Gerfen and Surmeier, 2011; McGregor and Nelson, 2019). One factor that is believed to contribute to this alteration are changes in striatal acetylcholine (ACh), arising mainly from local release by cholinergic interneurons (ChIs) (Kawaguchi, 1993; Poppi et al., 2021; Tanimura et al., 2018). As DA and ACh have multiple cooperative and reciprocal interactions within the striatum, the dynamic interplay between them is thought to be crucial for motor function (Aosaki et al., 1994; Avila et al., 2020; Howe et al., 2019; Krok et al., 2023; Straub et al., 2014; Threlfell et al., 2012), and their disruption contributing to movement impairment in PD (Aosaki et al., 2010; Barbeau, 1962; McGeer et al., 1961). In light of the importance of ACh for striatal function, both changes in ACh levels and ChI activity have been reported following DA loss (Aosaki et al., 1994; Choi et al., 2020; DeBoer et al., 1993; Ding et al., 2006; Lehmann and Langer, 1983; MacKenzie et al., 1989; Maurice et al., 2015; McKinley et al., 2019; Sanchez et al., 2011; Sanchez-Catasus et al., 2022; Spehlmann and Stahl, 1976), such that targeting the cholinergic system can improve motor deficits in PD and preclinical models (Fox et al., 2018; Katzenschlager et al., 2002; Kharkwal et al., 2016; Laverne et al., 2022; Maurice et al., 2015; Moehle et al., 2021; Ztaou et al., 2016). However, despite the critical role of ACh, it still remains poorly understood how cholinergic transmission itself and the modulation of striatal output are altered in PD.

ACh indirectly regulates MSN activity via muscarinic and nicotinic receptors on afferent inputs, as well as through direct modulation by postsynaptic muscarinic receptors. Muscarinic receptors are metabotropic G-protein coupled receptors (GPCRs) and while Gα_q_-coupled M1 receptors are expressed in all MSNs, Gα_i/o_-coupled M4 receptors are predominantly expressed in dMSNs (Bernard et al., 1992; Hersch et al., 1994; Levey et al., 1991; Yan et al., 2001). Due to this preferential location in the pathway critical for movement facilitation, M4 receptors have been regarded as the primary muscarinic subtype involved in DA-ACh interactions, reliably decoding ChI firing patterns (Mamaligas and Ford, 2016) and regulating DA transmission and associated motor behaviors (Moehle and Conn, 2019). M4 receptors are believed to participate in the control motor function by modulating the presynaptic release of DA (Foster et al., 2016; Jeon et al., 2010), and by postsynaptically opposing Gα_olf/s_-coupled D1-receptor signaling, which regulates the induction of corticostriatal synaptic plasticity (Moehle et al., 2017; Nair et al., 2019; Onali and Olianas, 2002; Shen et al., 2015). Together these findings emphasize the importance of M4-receptor signaling in the direct pathway for correct striatal function.

The long-held DA-ACh balance hypothesis assumes opposing roles for these neuromodulators, where DA depletion is accompanied by an increase in ACh such that PD is considered a hypercholinergic state (Aosaki et al., 2010; Barbeau, 1962; McGeer et al., 1961). In dMSNs, the expected changes in DA and ACh levels are predicted to lead to reduced D1-signaling and enhanced of M4-signaling, resulting in an overall inhibition of the direct pathway and movement. In order to counteract the loss of DA, the standard treatment in PD utilizes the precursor levodopa (L-DOPA). While this remains effective in early stages, eventual side effects occur including the onset of abnormal involuntary movements referred to as levodopa-induced dyskinesia (LID) (Cotzias et al., 1969). However, though an enhancement of M4-mediated cholinergic transmission is predicted by the DA-ACh hypothesis, directly testing these changes following DA loss and its replacement by L-DOPA has yet to be done. Here we examine the strength of cholinergic transmission across striatal regions and find that direct pathway M4-transmission is reduced in parkinsonian mice, which occurs as a result of downregulation of postsynaptic receptor function and parallels the loss of DA. As restoring aberrant M4-transmission partially rescued motor deficits and alleviated dyskinetic behaviors, these results suggest an underappreciated role for direct pathway M4-transmission in the pathogenesis of PD and the development of LID.

## RESULTS

### The strength of M4-mediated ACh transmission in dMSNs differs across the striatum

To examine M4-mediated cholinergic transmission in dMSNs an electrophysiological approach was used, where G-protein-coupled inwardly rectifying K^+^ channels (GIRK2; Kir3.2) and a tdTomato fluorophore were exogenously expressed, by unilaterally injecting AAV.DIO.GIRK2.T2A.tdTomato into the DSt of D1-Cre (*Drd1-Cre^+/-^*) mice (Figure 1A). As M4 receptors in dMSNs signal via second-messenger intracellular cascades that modulate postsynaptic conductances rather than directly activating them, overexpression of GIRK2 channels provides a *de novo* electrophysiological readout for M4 activation and signaling in these cells (Cai and Ford, 2018; Mamaligas et al., 2019; Mamaligas and Ford, 2016). GIRK2 expression does not have measurable effects on passive electrophysiological properties of MSNs (Marcott et al., 2014), apparent affinity of M4 receptors (Mamaligas and Ford, 2016) or behavioral performance in basic motor tasks (Gong et al., 2021).

**Figure 1:**
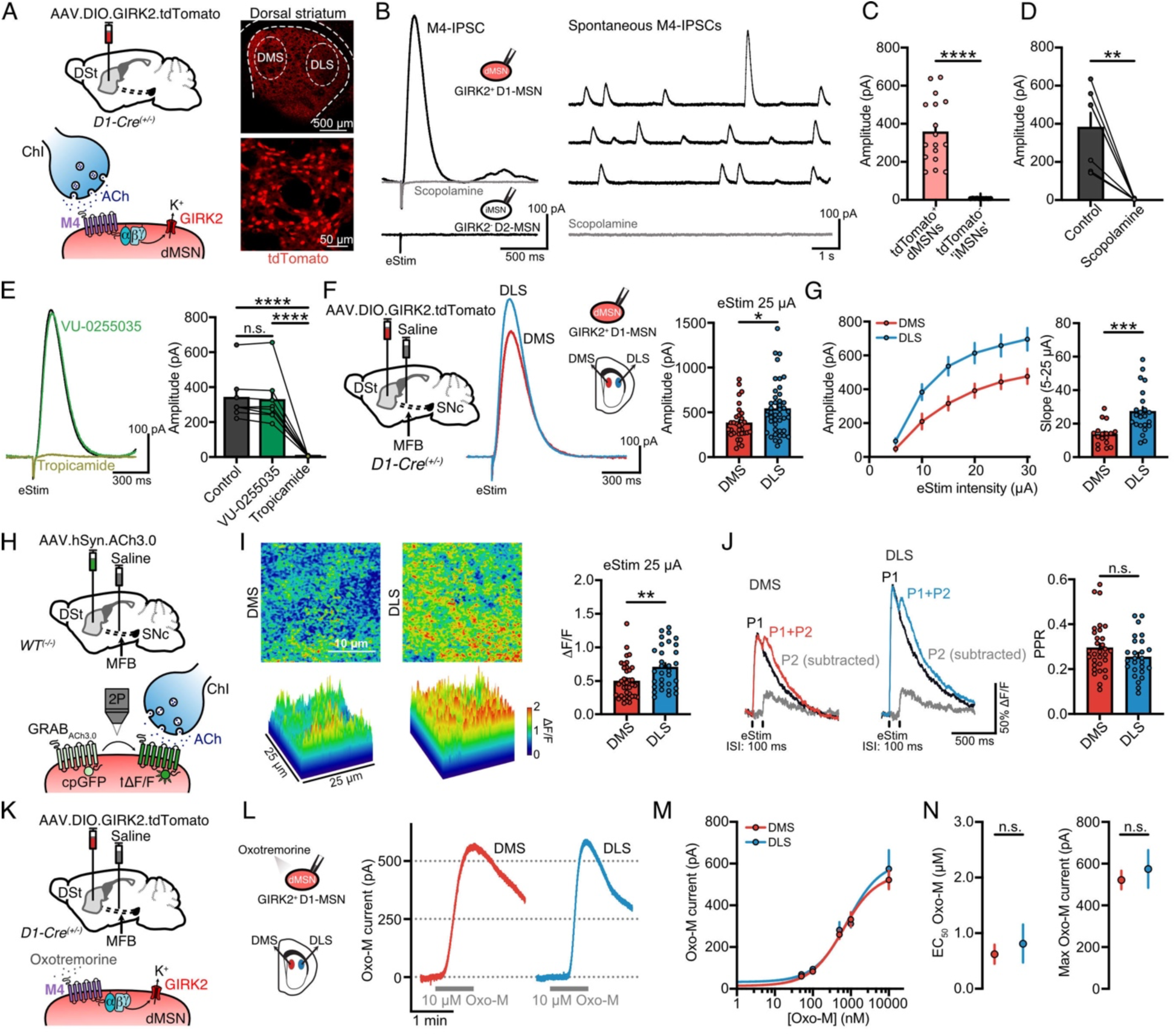
Differential M4-mediated cholinergic transmission in dMSNs across dorsal striatum regions. (A) Schematics of AAV9.hSyn.DIO.tdTomato.T2A.GIRK2 injection into the DSt of D1-Cre mice (top left) and ChI-dMSNs synapse showing the intracellular coupling between endogenous M4 receptors and exogenous GIRK2 overexpressed in dMSNs (bottom left). tdTomato fluorescence of a coronal striatal section highlighting DSt regions (top right) and close-up view of tdTomato^+^ individual dMSNs (bottom right). (B) Representative traces of electrically evoked (left) and spontaneous M4-IPSCs (right) recorded from tdTomato^+^ dMSNs (black) that are blocked by 1 µM scopolamine application (gray). No IPSCs are detected in tdTomato^-^ putative ‘iMSNs’ (bottom left). (C) Summary data of M4-IPSCs in tdTomato^+^ dMSNs and tdTomato^-^ putative ‘iMSNs’ (dMSNs: n=17, N=6; ‘iMSNs’: n=10, N=3; Mann-Whitney). (D) Quantification of scopolamine effect on M4-eIPSCs (n=7, N=3; paired t-test). (E) Representative traces and quantification of M4-eIPSCs following application of selective M1 (1µM VU-0255035) and M4 (1µM tropicamide) receptors antagonists (n=8, N=3; RM One-Way ANOVA, Holm-Šídák’s post-hoc test). (F) Schematics of AAV9.hSyn.DIO.tdTomato.T2A.GIRK2 and saline injections into the DSt and MFB respectively in D1-Cre mice (left). Representative traces and quantification of electrically evoked M4-IPSCs in DMS and DLS (25 µA, 0.5 ms) (right) (DMS: n=32, N=13; DLS: n=43, N=16; Mann-Whitney). (G) Plot of M4-IPSCs amplitudes versus electrical stimulation intensity for both striatal regions (left) and summary data of slope for 5-25 µA range (right) (DMS: n=17, N=9; DLS: n=23, N=12; Mann-Whitney). (H) Schematics of AAV.hSyn.ACh3.0 and saline injections into the DSt and MFB respectively in WT mice (top). Cartoon schematic showing the principle of GRAB_ACh3.0_ sensor function. (I) Representative fluorescence changes of GRAB_ACh3.0_ in DMS and DLS following a single electrical stimulus (25 µA, 0.5 ms) in the center of the square region of interest (25 µm x 25 µm), shown in 2-dimensions (top) and surface plot (bottom). Quantification of ΔF/F is shown on the bar chart (right) (DMS: n=38, N=9; DLS: n=32, N=9; Mann-Whitney). (J) Representative photometry traces of GRAB_ACh3.0_ fluorescence with single (P1) and paired-pulse stimulation (P1+P2, interstimulus interval (ISI): 100ms), and the digitally subtracted P2 component on gray. Summary data of PPR is shown on the bar chart (right) (DMS: n=31, N=7; DLS: n=26; N=7; Mann-Whitney). (K) Schematics of AAV9.hSyn.DIO.tdTomato.T2A.GIRK2 and saline injections into the DSt and MFB respectively in D1-Cre mice (top). Cartoon schematic showing the bath application of Oxo-M. (L) Representative traces of M4-mediated Oxo-M currents following bath application of a saturating concentration of Oxo-M (10 µM). Spontaneous M4-IPSCs and electrical artifacts were blanked for clarity. (M) Oxotremorine concentration-response curve for M4 receptors across striatal regions. (N) EC_50_ values from oxotremorine concentration-response curves in (M) (left; DMS: n=52, N=5-10; DLS: n=54, N=5-12; unpaired t-test) and maximal current values at 10 µM Oxo-M (right: DMS: n=14, N=10; DLS: n=12, N=10; Mann-Whitney). Summary data is mean ± SEM. n: number of cells, N: number of mice; n.s. p>0.05; *p<0.05; **p<0.01; ***p<0.001; ****p<0.0001. See also Table S1.

In the presence of antagonists to block glutamate, GABA, DA, and nicotinic receptors, the pacemaker firing of ChIs evokes scopolamine-sensitive (1 µM) spontaneous muscarinic M4-mediated inhibitory postsynaptic currents (M4-IPSC) in tdTomato^+^ GIRK2^+^ dMSNs, as well as larger evoked M4-IPSCs when electrical stimulation (25 µA, 0.5 ms) is used to drive synchronous release of ACh from multiple ChIs (Mamaligas and Ford, 2016) (Figure 1B-1D). As M4 receptors are preferentially expressed in dMSNs (Bernard et al., 1992; Hersch et al., 1994; Levey et al., 1991; Yan et al., 2001), overexpression of GIRK2 channels leads to M4-IPSCs only in dMSNs (Figure 1C and S1A). Electrically evoked M4-IPSCs in dMSNs were unaffected by the M1-antagonist, VU-0255035 (1 µM) but were abolished following application of the M4-antagonist, tropicamide (1 µM), confirming that synaptic responses were mediated by M4 receptors (Figure 1E). M4-IPSCs in dMSNs were the result of ACh release from ChIs, as a single action potential in a ChI led to a time-locked unitary M4-IPSC in a synaptically-coupled dMSN when paired recordings were conducted (Figure S1B).

DSt can be subdivided into the dorsomedial (DMS) and dorsolateral striatum (DLS), which differ in afferent and efferent connections, behaviorally-relevant circuitry, and disease associated dysfunctions. These differences partially rely on dopaminergic inputs that project from distinct but partially overlapping DA neuron populations (Chen et al., 2021; Lerner et al., 2015). While differences in dopaminergic regulation of ChIs across regions are known (Cai and Ford, 2018; Chuhma et al., 2014), it remains less clear whether the intrinsic strength of cholinergic transmission also differs across striatal regions. To address this for M4-transmission, we proceeded to record M4-IPSCs across DMS and DLS (Figure 1F-1G). Saline was injected into the medial forebrain bundle (MFB) in these mice to serve as controls for subsequent experiments. Across a range of electrical stimulations, the amplitude of M4-IPSCs evoked by a single electrical stimulus was greater in DLS compared to DMS (Figure 1F-1G), indicating a greater strength of signaling in the dorsolateral region.

To identify the mechanism underlying enhanced M4-transmission in the DLS, we first quantified electrically evoked release of ACh across striatal regions in WT mice using the genetically encoded optical sensor GRAB_ACh3.0_ (Jing et al., 2020) and 2-photon fluorescent imaging (Figure 1H). A single electrical stimulus evoked a greater peak change in GRAB_ACh3.0_ fluorescence (ΔF/F) in the DLS than the DMS (Figure 1I). As bath application of a saturating concentration of ACh (100 µM; together with 100 nM acetylcholinesterase (AChE) inhibitor ambenonium) induced similar fluorescence increases in both DMS and DLS, the regional difference in electrically evoked release likely was not due to differences in sensor expression (Figure S1D). The difference in release was also unlikely due to regional differences in the probability of ACh release, as the paired-pulse ratio (PPR) of two stimuli (100 ms inter-stimulus interval (ISI)), an indirect measure of release probability, was similar in the DMS and DLS when assayed with faster temporal imaging (Figure 1J). No difference was found in the postsynaptic sensitivity or efficacy of M4 receptors in dMSNs between the DMS and DLS when measuring GIRK2 currents, as concentration-response curves generated with the muscarinic agonist oxotremorine-M (Oxo-M) showed similar half-maximal effective concentration (EC_50_) and maximum outward currents across regions (Figure 1K-N). Thus, for a given stimulation, slightly more ACh is released in the DLS than in the DMS, which is in line with the faster activation kinetics of M4-IPSCs observed in dorsolateral dMSNs (Figure S1C) and may be the result of the higher density and clustering of ChIs (Matamales et al., 2016) or ChI arborizations that have been previously observed in the DLS (Burke and Karanas, 1990).

### M4-mediated cholinergic transmission is differentially reduced across striatal regions following partial or total DA depletion

We next examined how M4-transmission in dMSNs was altered following DA loss. To mimic early or advanced stages of the disease, we injected either a low (1 µg; LD) or high (4 µg; HD) dose of 6-hydroxydopamine (6-OHDA) unilaterally into the MFB to induce dopaminergic lesions resulting in either a partial or near complete depletion of striatal DA (Boix et al., 2015; Cai et al., 2021) (Figure 2A-2B). As stated above, control animals were injected with saline into the MFB, and the data from the previous results was used to compare control and 6-OHDA treated animals. Three to four weeks post 6-OHDA injection, tyrosine hydroxylase (TH) immunoreactivity was reduced in both the DSt and SNc (Figure 2B-2C). The partial lesion induced a ∼50-60% decrease in TH-immunoreactivity in the DSt and SNc (Figure 2B-2D), which correlates with the estimated ∼50% reduction in putamen DA fibers required to produce clinical motor symptoms in PD (Kordower et al., 2013), and is similar to the ∼57% reduction in evoked DA release we have previously described with low-dose 6-OHDA partial lesions (Cai et al., 2021). DA neurons within SNc differentially degenerate in PD, and the most vulnerable population is located in the ventral tier that projects to the putamen (Fearnley and Lees, 1991; Kish et al., 1988; Kordower et al., 2013). This is similar to animal lesion models of PD, where ventral tier SNc neurons and their DLS-projecting fibers are more sensitive to degeneration (Gerfen et al., 1987; Poulin et al., 2018). In agreement with this, low-dose 6-OHDA induced greater loss of TH-immunoreactivity in the DLS than DMS (Figure 2B, 2D). As expected, lesioning of the DA system resulted in motor behavioral deficits in 6-OHDA mice when assayed in either the cylinder or rotarod tests, with the cylinder test revealing asymmetry in forelimb use and the rotarod test revealing decreased latency to fall (Figure 2E-2F). The extent of both motor deficits was correlated with the loss of TH^+^ fibers in the DSt (Figure 2E-2F). The presence of motor deficits in the cylinder test was subsequently used to include mice in their appropriate experimental groups, with a contralateral paw use below 40% being required to include mice in the complete DA depletion group (Tanimura et al., 2019), while contralateral paw use below 45% was chosen for including animals in the partial DA loss group (Figure 2E).

**Figure 2:**
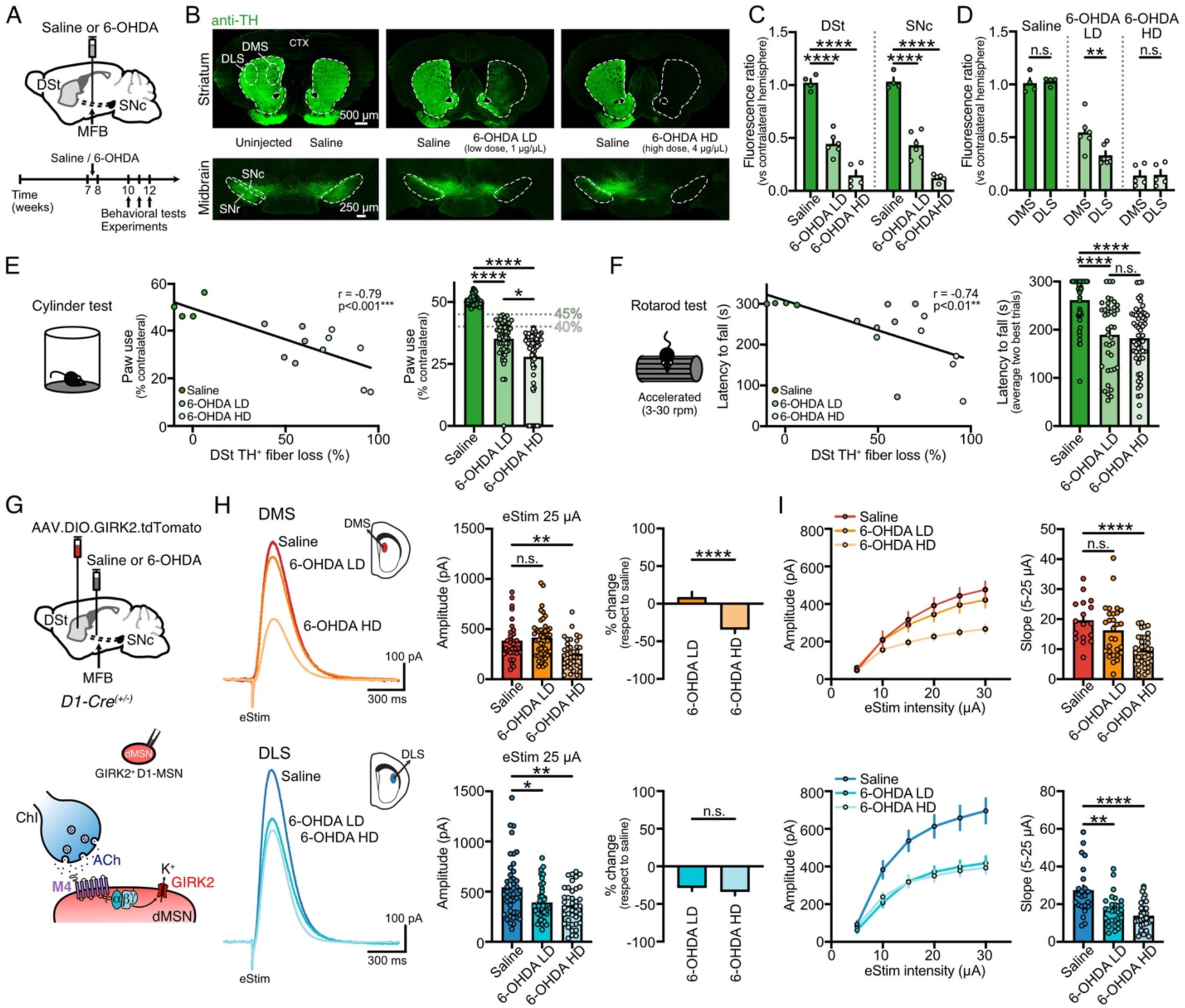
Partial and complete DA depletion differentially reduces M4-mediated cholinergic transmission across dorsal striatum regions. (A) Schematics of saline or 6-OHDA injections into the MFB (top) and timeline for experiments (bottom). (B) Striatal and midbrain coronal sections showing TH-immunoreactivity following unilateral saline, 6-OHDA LD (1 µg/µL) or 6-OHDA HD (4 µg/µL) injections. (C) Quantification of TH-immunoreactivity in DSt and SNc expressed as the fluorescence ratio of the injected hemisphere versus the uninjected/saline-injected hemisphere (saline: N=4, 6-OHDA LD: N=6, 6-OHDA HD: N=5; One-Way ANOVA, Dunnet’s post-hoc test). (D) Quantification of TH-immunoreactivity across DMS and DLS for all conditions (saline: N=4, Mann-Whitney; 6-OHDA LD: N=6, Mann-Whitney; 6-OHDA HD: N=5, unpaired t-test). (E) Cylinder test performance and percent of DSt TH^+^ fiber loss correlation (left) (n=15; Pearson correlation). Summary data of contralateral paw use in cylinder test for all D1-Cre and WT mice included in this study (right). Cutoff values for 6-OHDA LD and 6-OHDA HD shown on the bar chart were used as inclusion criteria for partial and total DA loss experimental groups (saline: N=60, 6-OHDA LD: N=56; 6-OHDA HD: N=63; Kruskal-Wallis, Dunn’s post-hoc test). (F) Rotarod test performance and percent of DSt TH^+^ fiber loss correlation (left) (n=15; Spearman correlation). Summary data of latency to fall for all conditions (saline: N=42, 6-OHDA LD: N=39; 6-OHDA HD: N=57; Kruskal-Wallis, Dunn’s post-hoc test). (G) Schematics of GIRK2-encoding Cre-dependent virus and saline/6-OHDA injections into DSt and MFB respectively in D1-Cre mice (top) and ChI-dMSNs synapse showing the intracellular coupling between M4 and GIRK2 in dMSNs (bottom). (H) Representative traces and quantification of electrically evoked M4-IPSCs (25 µA, 0.5 ms) in DMS (top) and DLS (bottom) for saline (control data taken from Figure 1F), 6-OHDA LD and 6-OHDA HD conditions (DMS: saline n=32, N=13; 6-OHDA LD n=45, N=12; 6-OHDA HD n=37, N=11; Kruskal-Wallis, Dunn’s post-hoc test / DLS: saline n=43, N=16; 6-OHDA LD n=44, N=15; 6-OHDA HD n=39, N=13; Kruskal-Wallis, Dunn’s post-hoc test). Percent change quantification of M4-IPSCs following partial and complete DA loss with respect to saline is shown on the right (Mann-Whitney). (I) Plot of M4-IPSCs amplitudes versus electrical stimulation intensity following DA loss (left) and summary data of slope for 5-25 µA range (right) for DMS (top) and DLS (bottom). Control data for saline condition taken from Figure 1G (DMS: saline n=17, N=9; 6-OHDA LD n=29, N=9; 6-OHDA HD n=36, N=11; One-Way ANOVA, Dunnet’s post-hoc test / DLS: saline n=23, N=12; 6-OHDA LD n=29, N=11; 6-OHDA HD n=35, N=12; One-Way ANOVA, Dunnet’s post-hoc test). Summary data is mean ± SEM. n: number of cells, N: number of mice; n.s. p>0.05; *p<0.05; **p<0.01; ***p<0.001; ****p<0.0001. See also Table S1.

Partial DA loss selectively reduced electrically evoked M4-mediated transmission in dMSNs in the DLS, but not DMS (Figure 2G-2H). The reduction in M4-IPSCs paralleled the extent of dopaminergic lesion noted above (Figure 2B, 2D) suggesting it was driven by the gradual loss of DA. A high dose of 6-OHDA, which drives a more complete loss of striatal dopaminergic innervation, led to a decrease in M4-IPSCs in the DMS but had no further effect on M4-transmission in the DLS compared to a partial lesion (Figure 2H). Similar results could be seen over a range of electrical stimuli examining input/output relationships across conditions (Figure 2I). Thus, while PD is often thought to exhibit enhanced cholinergic transmission (Barbeau, 1962; Lehmann and Langer, 1983; McGeer et al., 1961; Spehlmann and Stahl, 1976), these results indicate that direct pathway M4-mediated cholinergic transmission is reduced following DA loss.

### Reduced M4-mediated cholinergic transmission in response to DA loss is due to decreased postsynaptic M4-function and not presynaptic ACh release

We next set out to examine the mechanisms that underlie the decrease in strength of M4-transmission in dMSNs following DA loss. We first examined potential presynaptic changes in electrically evoked ACh release, using GRAB_ACh3.0_ to quantify changes in 6-OHDA lesioned parkinsonian mice (Figure 3A) under similar experimental conditions as above. In the DMS, ACh release was unaffected by low-dose 6-OHDA treatment but was increased when striatal DA was completely depleted (Figure 3B). This was in contrast to the DLS, where ACh release did not exhibit any significant alterations following either partial or total DA loss (Figure 3B). In both regions, the PPR following two stimulations (100 ms ISI) was unaffected by either low- or high-dose 6-OHDA treatment, suggesting that the probability of electrically evoked ACh release is not altered following DA loss in either DMS or DLS (Figure 3C). Thus, the reduction in M4-transmission was not due to changes in ACh release, since it remained unaffected in DLS, the most sensitive region to dopaminergic denervation, and increased only in DMS when DA was nearly completely depleted.

**Figure 3:**
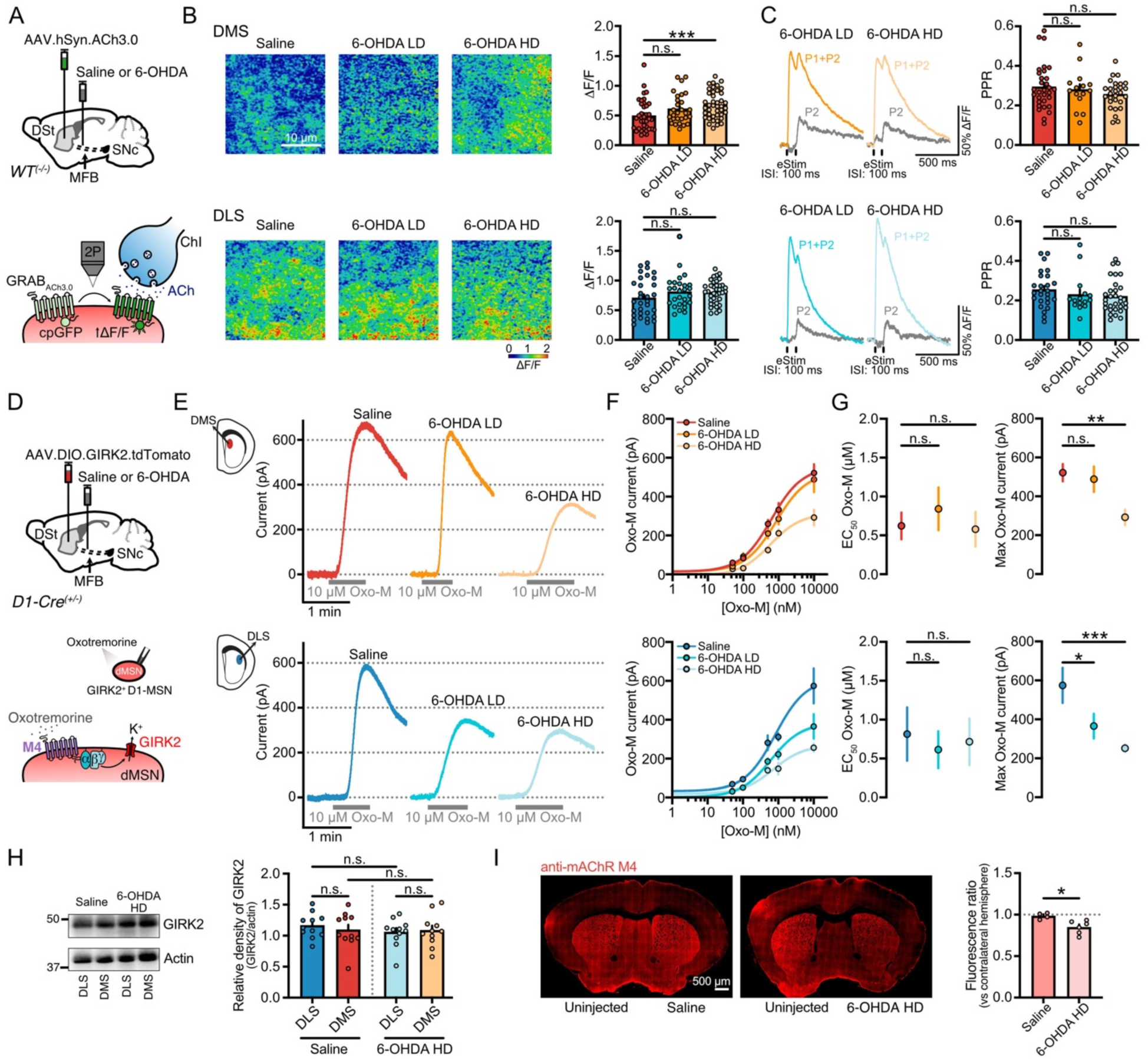
DA depletion reduces M4-transmission through a postsynaptic mechanism and not as a result of changes in presynaptic ACh release. (A) Schematics of AAV.hSyn.ACh3.0 and saline/6-OHDA injections into the DSt and MFB respectively in WT mice (top). Cartoon schematic showing the principle of GRAB_ACh3.0_ sensor function (bottom). (B) Representative fluorescence changes of GRAB_ACh3.0_ in DMS (top) and DLS (bottom) evoked by single electrical stimulation (25 µA, 0.5 ms) in the center of the square region of interest (25 µm x 25 µm) for saline, 6-OHDA LD and 6-OHDA HD conditions. Quantification of ΔF/F is shown on the bar charts (right). Control data for saline group taken from Figure 1I (DMS: saline n=38, N=9; 6-OHDA LD n=28, N=6; 6-OHDA HD n=42, N=9; Kruskal-Wallis, Dunn’s post-hoc test / DLS: saline n=32, N=9; 6-OHDA LD n=27, N=6; 6-OHDA HD n=38, N=9; Kruskal-Wallis). (C) Representative photometry traces of GRAB_ACh3.0_ fluorescence in DMS (top) and DLS (bottom) following DA loss with single (P1) and paired-pulse stimulation (P1+P2), and the digitally subtracted P2 component on gray. Summary data for PPR is shown on the bar charts (right). Control data for saline group taken from Figure 1J (DMS: saline n=31, N=7; 6-OHDA LD n=15, N=4; 6-OHDA HD n=28, N=5; Kruskal-Wallis / DLS: saline n=26, N=7; 6-OHDA LD n=15, N=4; 6-OHDA HD n=30, N=5; Kruskal-Wallis). (D) Schematics of AAV9.hSyn.DIO.tdTomato.T2A.GIRK2 and saline/6-OHDA injections into the DSt and MFB respectively in D1-Cre mice (top). Cartoon schematic showing the bath application of Oxo-M. (E) Representative traces of M4-mediated Oxo-M currents in DMS (top) and DLS (bottom) for saline (DLS saline taken from Figure 1L), 6-OHDA LD and 6-OHDA HD conditions following bath application of a saturating concentration of Oxo-M (10 µM). Spontaneous M4-IPSCs and electrical artifacts were blanked for clarity. (F) Oxotremorine concentration-response curve for M4 receptors in DMS (top) and DLS (bottom) for saline (taken from Figure 1M), 6-OHDA LD and 6-OHDA HD. (G) EC_50_ values from oxotremorine concentration-response curves in (F) (top left; DMS: saline n=52, N=5-10; 6-OHDA LD n=43, N=3-8; 6-OHDA HD n=59, N=5-11; One-Way ANOVA / bottom left; DLS: n=54, N=5-12; 6-OHDA LD n=64, N=6-12; 6-OHDA HD n=42, N=5-9; One-Way ANOVA) and maximal Oxo-M current values at 10 µM oxotremorine (top right: DMS: saline n=14, N=10; 6-OHDA LD n=8, N=6; 6-OHDA HD n=14, N=11; One-Way ANOVA, Dunnet’s post-hoc test / bottom right: DLS: saline n=12, N=10; 6-OHDA LD n=12, N=10; 6-OHDA HD n=11, N=9; Kruskal-Wallis, Dunn’s post-hoc test). (H) Representative western blot showing GIRK2 protein overexpression across DMS and DLS in saline and 6-OHDA HD conditions. Quantification of expression levels normalized to actin is shown on the right (N=11 for each condition; One-Way ANOVA). (I) Striatal coronal sections showing M4-immunoreactivity following unilateral injections of saline and 6-OHDA HD (left). Quantification of fluorescence ratio of the injected versus contralateral uninjected hemisphere (right) (saline N=4, 6-OHDA HD N=6; unpaired t-test). Summary data is mean ± SEM. n: number of cells, N: number of mice; n.s. p>0.05; *p<0.05; **p<0.01; ***p<0.001; ****p<0.0001. See also Table S1.

As striatal ACh is also regulated by AChE, we tested if ACh clearance was altered following DA loss by applying the AChE inhibitor ambenonium (10 nM) while recording electrically-evoked M4-IPSCs in GIRK2^+^ dMSNs. Because AChE limits the duration of M4-mediated transmission in dMSNs (Mamaligas and Ford, 2016), the net charge transfer and decay time constant of evoked synaptic currents were increased following AChE inhibition (Figure S2A-S2C). However, there was no difference in the effect of ambenonium across either the DMS or DLS following DA loss (Figure S2A-S2C). Together, these results suggest that changes in evoked ACh release from ChIs and/or its clearance likely does not explain the reduction in M4-mediated cholinergic transmission in dMSNs that occurs in response to striatal dopaminergic lesion.

Next, we investigated whether DA loss altered postsynaptic M4-function by bath applying the muscarinic agonist Oxo-M and generating concentration-response curves from M4-mediated outward currents in dMSNs (Figure 3D-3G). While there were no differences in M4 receptor sensitivity (EC_50_) across striatal regions following DA depletion, there was a significant reduction in efficacy as measured by the maximum outward current evoked by a saturating concentration of Oxo-M (10 µM) (Figure 3E-3G). In the DMS, maximum outward currents were decreased only with the complete loss of DA and were unaffected following partial DA depletion (Figure 3F-3G). In contrast, maximum outward currents were significantly decreased in the DLS by both partial and complete dopaminergic lesions (Figure 3F-3G). The 6-OHDA-induced decrease in M4-mediated currents was not due to differences in GIRK2 expression, as GIRK2 protein levels were similar across regions in all conditions (Figure 3H). Instead, the reduction in M4-function may be at least partially the result of a decrease in M4 receptor expression, as immunostaining against M4 showed that unilaterally injected high-dose 6-OHDA treated mice had a ∼20% reduction in total M4-immunoreactivity across the DSt compared to the saline-injected controls (Figure 3I). While M4 receptors in the striatum are also expressed in ChIs and corticothalamic terminals, they are most highly expressed in dMSNs (Hersch et al., 1994; Levey et al., 1991), suggesting that at least some of the decrease in striatal M4-immunoreactivity results from a decrease in receptor levels in dMSNs. Together, these results indicate that the region-specific reduction in the amplitude of evoked M4-IPSCs following DA loss is due to a decrease in postsynaptic M4-function.

As we found that M4 receptor protein levels were reduced by ∼20% but transmission and postsynaptic M4-function were reduced by ∼35-55%, additional alterations in downstream signaling cascades may occur. In an attempt to rescue the impaired M4-mediated cholinergic transmission, and to further dissect the contribution of each potential mechanism, we next developed two different strategies to either increase M4 receptor expression or enhance downstream M4-signaling selectively in dMSNs.

### Overexpression of M4 receptors in dMSNs restores M4-transmission and ameliorates balance and coordination motor deficits

We designed an AAV encoding for the Cre-dependent expression of the M4 receptor and a fluorescent reporter eGFP to test whether selective overexpression in dMSNs could rescue deficits in cholinergic transmission in parkinsonian mice. We first validated this construct in control saline-treated mice. Three weeks following injection of AAV.DIO.M4.eGFP into the DSt of D1-Cre mice, M4-immunoreactivity was increased by ∼60% in the injected hemisphere (Figure 4A). Co-injection of AAV.DIO.M4.eGFP together with AAV.DIO.GIRK2.T2A.tdTomato in D1-Cre mice led to widespread eGFP and tdTomato fluorescence in overlapping populations of neurons (Figure 4B). Recordings from eGFP^+^/tdTomato^+^ dMSNs revealed that the amplitude of both M4-IPSCs (Figure 4C), and M4-mediated outward currents evoked by Oxo-M (10 µM) (Figure 4D) were greater in M4-overexpressing mice (M4 OE) than in control mice (eYFP), in which AAV.DIO.GIRK2.T2A.tdTomato was co-injected with a control fluorophore (AAV.DIO.eYFP) (Figure S3A). Thus, overexpressed M4 receptors were functional and coupled to downstream effectors to mediate postsynaptic responses in dMSNs.

**Figure 4:**
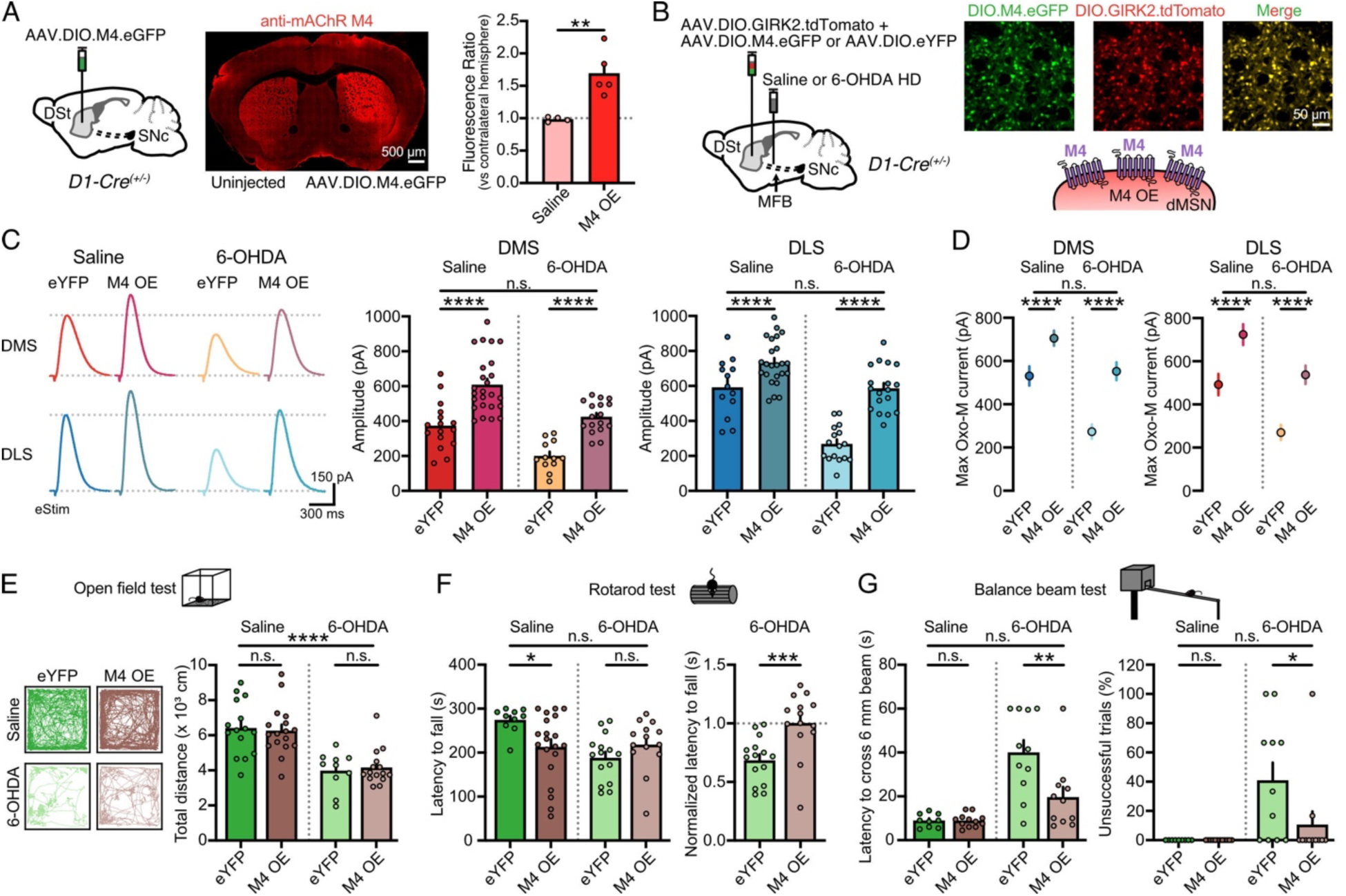
Overexpression of M4 receptors in dMSNs restores M4-transmission and ameliorates motor deficits in parkinsonian mice. (A) Schematics of AAV.DIO.M4.eGFP unilateral injection into the DSt of D1-Cre mice, and striatal coronal section showing M4-immunoreactivity (left). Quantification of fluorescence ratio of the injected versus contralateral uninjected hemisphere (right). Saline data taken from Figure 3I (saline N=4, M4 OE N=5; unpaired t-test). (B) Schematics of AAV.DIO.M4.eGFP or AAV.DIO.eYFP co-injections with AAV9.hSyn.DIO.tdTomato.T2A.GIRK2 into the DSt, and saline/6-OHDA HD injections into the MFB of D1-Cre mice (left). Close-up images of dMSNs co-expressing eGFP and tdTomato fluorescent reporters and cartoon schematic of a dMSN depicting M4 OE (right). (C) Representative traces (left) and quantification (right) of electrically evoked M4-IPSCs (25 µA, 0.5 ms) in DMS and DLS for eYFP saline, M4 OE saline, eYFP 6-OHDA and M4 OE 6-OHDA conditions (DMS: eYFP saline: n=16, N=6; M4 OE saline: n=25, N=9; eYFP 6-OHDA: n=12, N=5; M4 OE 6-OHDA: n=17, N=7 / DLS: eYFP saline: n=13, N=6; M4 OE saline: n=24, N=8; eYFP 6-OHDA: n=16, N=5; M4 OE 6-OHDA: n=18, N=7; Two-Way ANOVA, Šídák’s post-hoc test). (D) Summary data of maximal M4-mediated Oxo-M currents (10 µM) in DMS (left) and DLS (right) for eYFP saline, M4 OE saline, eYFP 6-OHDA and M4 OE 6-OHDA conditions (DMS: eYFP saline: n=12, N=6; M4 OE saline: n=16, N=9; eYFP 6-OHDA: n=9, N=5; M4 OE 6-OHDA: n=9, N=5 / DLS: eYFP saline: n=13, N=6; M4 OE saline: n=20, N=10; eYFP 6-OHDA: n=10, N=5; M4 OE 6-OHDA: n=9, N=5; Two-Way ANOVA, Šídák’s post-hoc test). (E) Example traces of locomotor activity in the open field test and quantification of total distance travelled (eYFP saline: N=15; M4 OE saline: N=17; eYFP 6-OHDA: N=10; M4 OE 6-OHDA: N=15; Two-Way ANOVA, Šídák’s post-hoc test). (F) Summary data of rotarod performance (latency to fall) shown as absolute (left) and normalized values (right) (eYFP saline: N=10; M4 OE saline: N=20; eYFP 6-OHDA: N=15; M4 OE 6-OHDA: N=14; Two-Way ANOVA, Šídák’s post-hoc test for absolute data and Mann-Whitney for normalized values). (G) Summary data of latency to cross 6 mm beam (left) and percentage of unsuccessful trials (right) in balance beam test (eYFP saline: N=9; M4 OE saline: N=12; eYFP 6-OHDA: N=11; M4 OE 6-OHDA: N=11; Two-Way ANOVA, Šídák’s post-hoc test). Summary data is mean ± SEM. n: number of cells, N: number of mice; n.s. p>0.05; *p<0.05; **p<0.01; ***p<0.001; ****p<0.0001. See also Table S1.

We next tested whether selective M4-overexpression in dMSNs could limit the decrease in cholinergic transmission following complete DA loss. In agreement with our previous results, the amplitude of M4-IPSCs of control eYFP mice was reduced across both the DMS and DLS following high-dose 6-OHDA treatment (Figure 4C). This deficit, however, was rescued in high-dose 6-OHDA-treated mice overexpressing M4 receptors such that despite complete DA depletion, the amplitude of M4-IPSCs was similar to saline-treated eYFP controls (Figure 4C). The rescue of M4-transmission in parkinsonian mice could be seen across a range of electrical stimulation intensities (Figure S3B-S3C) and also when examining postsynaptic M4-mediated outward currents evoked by bath application of Oxo-M (10 µM) (Figure 4D). Thus, selective overexpression of M4 receptors in dMSNs was sufficient to restore the impairment in M4-mediated cholinergic transmission that followed DA depletion back to normal levels.

While motor deficits like bradykinesia and rigidity can be effectively rescued by DA-replacement therapy in PD and preclinical models, certain motor features associated with impaired balance and coordination, such as postural instability, gait dysfunction, freezing of gait and falls, can be more refractory, suggesting the involvement of non-dopaminergic signaling pathways (McKay et al., 2019; Sethi, 2008; Wenger et al., 2022). To examine this, we next looked at the impact of restoring direct pathway cholinergic transmission on motor behavior by overexpressing M4 receptors in dMSNs in parkinsonian mice. We found that M4-overexpression had no effect on spontaneous locomotor activity (distance traveled in an open field) in control or 6-OHDA treated mice (Figure 4E), but it improved motor performance in the rotarod and balance beam tests in parkinsonian mice (Figure 4F-4G). As overexpression of M4 receptors led to a basal impairment in the rotarod task (Figure 4F), which was likely due to over-inhibition of the direct pathway (Moehle et al., 2017; Moehle and Conn, 2019), the motor improvement was revealed only after normalizing the latency to fall (Figure 4F). In contrast, in the balance beam test, M4-overexpression did not have any effect on basal performance so that the rescue in the 6-OHDA-induced deficits could be directly seen as a reduction in the latency to cross the beam and a reduction in foot slips despite the complete DA lesion (Figures 4G and S3F-S3G).

DA depletion also reduces engagement in walking tasks, and parkinsonian mice often remain immobile or stop while attempting to cross the beam, thus failing to complete a balance beam trial. The ‘start hesitation’ behavior has been described in rodents as being similar to the ‘freezing of gait’ observed in PD patients, a motor symptom often resistant to DA-replacement therapy (McKay et al., 2019; Sethi, 2008; Wenger et al., 2022) that may be related to brain cholinergic systems dysfunction (Avila et al., 2020; Bohnen et al., 2022, 2021; Pasquini et al., 2021; Xiao et al., 2017). To examine this, we also quantified the number of unsuccessful trials that failed due to freezing, out of the total number of sessions (Xiao et al., 2017). As opposed to high-dose 6-OHDA treated mice which frequently froze, M4-overexpressing mice showed reduced hesitation, such that they succeeded in crossing the beam as frequently as eYFP-expressing saline control mice (Figure 4G). Together, these results indicate that selective M4-overexpression in dMSNs restores the 6-OHDA-induced reduction in M4-mediated cholinergic transmission and can improve balance and coordination performance without further impairing locomotion, thus rescuing aspects of motor function that are often less responsive to standard DA-replacement therapy in PD.

### Selective ablation of RGS4 from dMSNs rescues M4-transmission and alleviates parkinsonian motor deficits

Next, we tested how manipulating M4 receptor intracellular regulatory cascades could alter cholinergic signaling and motor behavior in high-dose 6-OHDA treated mice. The timing and extent of GPCR signaling is controlled by several classes of GTPase-activating proteins (GAP) such as regulator of G-protein signaling (RGS) proteins that accelerate GTP hydrolysis and facilitate termination of G-protein signaling (Ross and Wilkie, 2000). In the striatum, RGS4 is highly expressed, regulates Gα_q_ and Gα_i/o_ (yet is biased towards Gα_i/o_) (Berman et al., 1996; Masuho et al., 2020) and has been implicated in the regulation of cholinergic and dopaminergic signaling, being relevant in several neurological disorders, including PD (Ahlers-Dannen et al., 2020; Ding et al., 2006; Geurts et al., 2003; Lerner and Kreitzer, 2012; Shen et al., 2015; Taymans et al., 2004). As RGS4 preferentially regulates signaling by Gα_i/o_-coupled GPCRs such as M4 receptors, we reasoned that selective ablation of RGS4 may enhance M4-signaling and thus rescue the reduction in M4-transmission which arises following DA depletion (Figure 5A-5B). To test this, we generated RGS4 conditional knockout mice (RGS4 cKO) (*RGS4^fl/fl^; D1-Cre^+/-^*), by crossing RGS4-floxed (Avrampou et al., 2019; Han et al., 2010; Stratinaki et al., 2013) and D1-Cre mice (Figure 5A). Knockout of RGS4 increased the amplitude of electrically evoked M4-IPSCs (Figures S4A and 5C) and prolonged the kinetics of activation and deactivation compared to littermate controls (*RGS4^wt/wt^; D1-Cre^+/-^*) (Figure S4A), confirming that RGS4 regulates M4-signaling in dMSNs. Maximum outward currents evoked by Oxo-M (10 µM) were also increased, while M4 sensitivity remained unaffected (Figure 5D and S4B).

**Figure 5:**
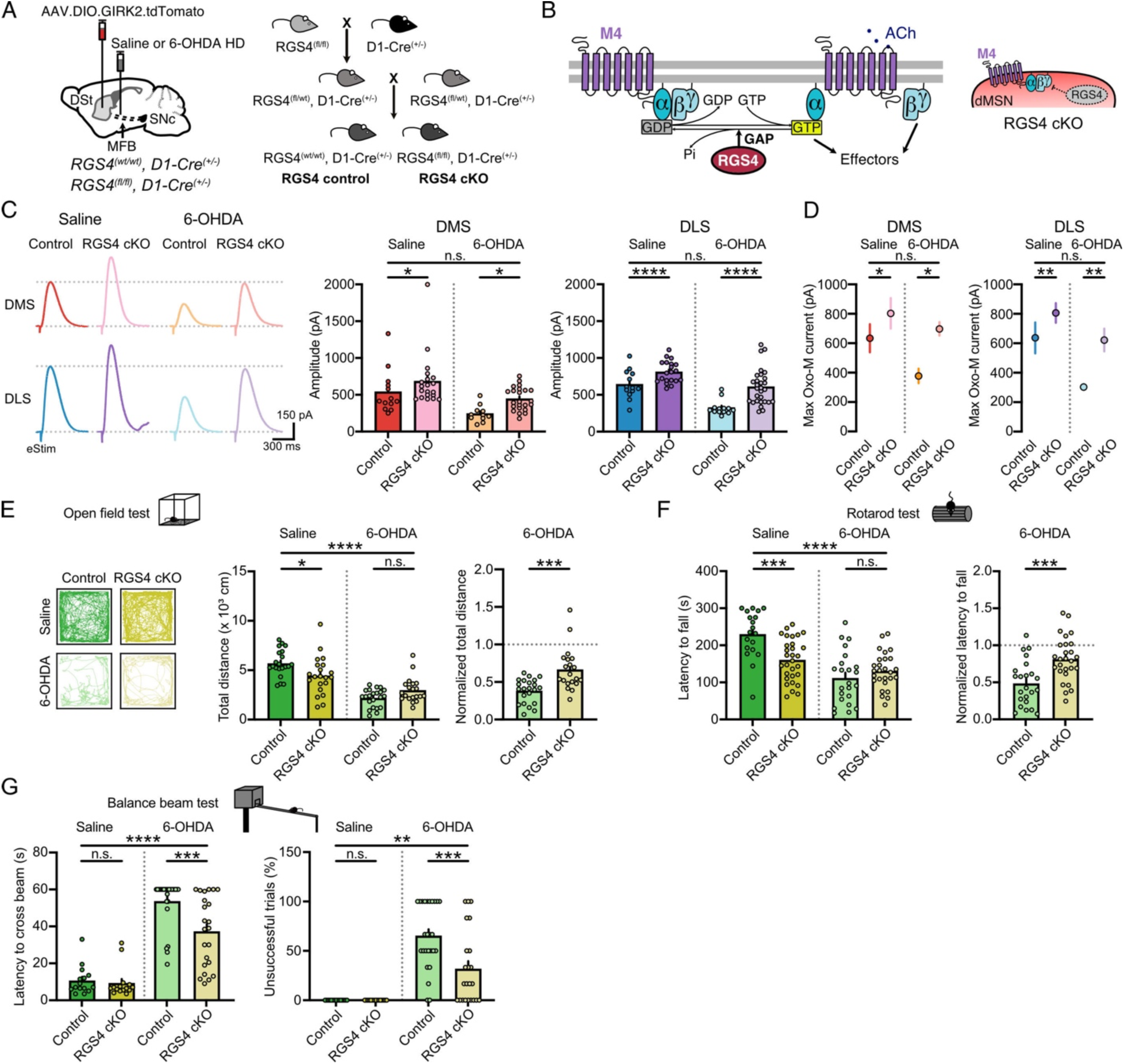
Selective ablation of RGS4 in dMSNs rescues M4-transmission and alleviates motor impairment in parkinsonian mice. (A) Schematics of AAV9.hSyn.DIO.tdTomato.T2A.GIRK2 and saline/6-OHDA HD injections into the DSt and MFB respectively in RGS4 control and RGS4 cKO mice (left). Generation of RGS4 cKO in dMSNs (RGS4^fl/fl^, D1-Cre^+/-^) and RGS4 control littermates (RGS4^wt/wt^, D1-Cre^+/-^) (right). (B) Schematic showing the effect of RGS4 on G-protein signaling for M4 receptors (left). Cartoon illustrating the ablation of RGS4 protein in dMSNs in RGS4 cKO strategy (right). (C) Representative traces (left) and quantification (right) of electrically evoked M4-IPSCs (25 µA, 0.5 ms) in DMS and DLS for control saline, RGS4 cKO saline, control 6-OHDA and RGS4 cKO 6-OHDA conditions (DMS: control saline: n=13, N=6; RGS4 cKO saline: n=20, N=11; control 6-OHDA: n=11, N=6; RGS4 cKO 6-OHDA: n=23, N=11 / DLS: control saline: n=13, N=7; RGS4 cKO saline: n=24, N=11; control 6-OHDA: n=13, N=6; RGS4 cKO 6-OHDA: n=29, N=10; Two-Way ANOVA, Šídák’s post-hoc test). (D) Summary data of maximal M4-mediated Oxo-M currents (10 µM) in DMS (left) and DLS (right) for control saline, RGS4 cKO saline, control 6-OHDA and RGS4 cKO 6-OHDA conditions (DMS: control saline: n=6, N=5; RGS4 cKO saline: n=15, N=10; control 6-OHDA: n=7, N=6; RGS4 cKO 6-OHDA: n=9, N=6 / DLS: control saline: n=6, N=5; RGS4 cKO saline: n=15, N=10; control 6-OHDA: n=8, N=5; RGS4 cKO 6-OHDA: n=9, N=7; Two-Way ANOVA, Šídák’s post-hoc test). (E) Example traces of locomotor activity in the open field test with quantification of total distance traveled in absolute (left) and normalized values (right) (control saline: N=21; RGS4 cKO saline: N=21; control 6-OHDA: N=22; RGS4 cKO 6-OHDA: N=20; Two-Way ANOVA, Šídák’s post-hoc test for absolute data and Mann-Whitney for normalized values). (F) Summary data of rotarod performance (latency to fall) shown as absolute (left) and normalized values (right) (control saline: N=19; RGS4 cKO saline: N=30; control 6-OHDA: N=23; RGS4 cKO 6-OHDA: N=27; Two-Way ANOVA, Šídák’s post-hoc test for absolute data and unpaired t-test for normalized values). (G)Summary data of latency to cross 6 mm beam (left) and percentage of unsuccessful trials (right) in balance beam test (control saline: N=15; RGS4 cKO saline: N=15; control 6-OHDA: N=26; RGS4 cKO 6-OHDA: N=23; Two-Way ANOVA, Šídák’s post-hoc test). Summary data is mean ± SEM. n: number of cells, N: number of mice; n.s. p>0.05; *p<0.05; **p<0.01; ***p<0.001; ****p<0.0001. See also Table S1.

To determine if prolonging M4-signaling by selective removal of RGS4 might limit the reduction in cholinergic transmission following DA depletion, we again performed a complete DA lesion using a high-dose of 6-OHDA (Figure 5A). In line with our previous findings, the amplitude of M4-IPSCs in control mice was decreased across DSt regions in response to DA loss (Figure 5C). Remarkably, the impairment in cholinergic transmission was restored in high-dose 6-OHDA mice selectively lacking RGS4 in dMSN, such that despite DA depletion, the amplitude of evoked M4-IPSCs was undistinguishable from saline-treated controls (Figure 5C). This rescue of M4-transmission in parkinsonian mice was observed across a range of electrical stimulation intensities (Figure S4C) and also when examining maximal postsynaptic M4-mediated outward currents evoked by bath application of Oxo-M (10 µM) (Figure 5D). Thus, selective ablation of RGS4 in dMSNs was sufficient to rescue the impairment in M4-mediated cholinergic transmission following DA lesion, restoring levels of transmission back to those of control.

We next examined the impact of selective RGS4 removal in dMSNs on parkinsonian motor behavior. While global knockout of RGS4 has been shown to either decrease (Lerner and Kreitzer, 2012) or increase (Ashrafi et al., 2017) 6-OHDA-induced motor impairments, cell-type specific effects of RGS4 ablation from dMSNs have not been addressed. Both rotarod performance and basal open field locomotor activity were slightly impaired in RGS4 cKO mice (Figure 5E-5F). As the extent of M4-transmission was enhanced to similar levels by either overexpressing M4 or prolonging its intracellular signaling (Figures 4C and 5C), the additional impairment in basal locomotion might be due to alterations in RGS4-mediated regulation of other Gα_i/o_- or Gα_q_-coupled GPCRs in dMSNs or other D1-expressing cells. Normalizing activity to account for these basal impairments, however, revealed a rescue of both locomotion (Figure 5E) and latency to fall from rotarod (Figure 5F) following RGS4 ablation in high-dose 6-OHDA-treated parkinsonian mice. Conditional knockout of RGS4 in dMSNs had no effect on basal performance in the balance beam test, but again led to a partial rescue of 6-OHDA-induced motor deficits including improvements in the latency to cross the beam (Figures 5G and S4F), reductions in the number of foot slips (Figure S4G) and reductions in ‘start hesitation’ or ‘freezing of gait’ episodes (Figure 5G). Overall, our results demonstrate that selective ablation of RGS4 from dMSNs restores the reduced M4-mediated cholinergic transmission and postsynaptic M4-function in response to DA lesion and partially rescues spontaneous locomotion, balance, and coordination deficits in parkinsonian mice. Hence, restoration of M4-mediated cholinergic transmission might be relevant for novel therapeutic approaches that complement standard L-DOPA treatment.

### Restoration of M4-transmission alleviates levodopa-induced dyskinesia

As PD progresses, L-DOPA administration eventually leads to the development of levodopa-induced dyskinetic involuntary movements (Bandopadhyay et al., 2022; Cotzias et al., 1969; Kwon et al., 2022). Although LID pathogenesis remains incompletely understood, the combination of dopaminergic denervation and chronic pulsatile stimulation of DA receptors is thought to drive an enhancement of the direct pathway due to activation of supersensitive D1-receptors and their downstream signaling cascade (Figure 6A) (Feyder et al., 2011; Gerfen, 2003; Girasole et al., 2018; Kwon et al., 2022; Parker et al., 2018; Picconi et al., 2003; Ryan et al., 2018; Spigolon and Fisone, 2018). In dMSNs, Gα_i/o_-coupled M4 receptors directly oppose Gα_olf_-coupled D1-receptor signaling (Figure 6A) (Moehle et al., 2017; Nair et al., 2019; Onali and Olianas, 2002; Shen et al., 2015). Historically as PD has been assumed as a hypercholinergic state (Aosaki et al., 2010; Barbeau, 1962; McGeer et al., 1961), the combination of low DA and high ACh tone has been thought to result in a strong inhibition of the direct pathway and suppression of movement through reduced D1-signaling and enhanced M4-signaling in dMSNs. Following from this, L-DOPA treatment is thought to rescue the reduced D1-signaling, with the heightened D1-receptor supersensitivity potentially balanced by an enhanced M4-signaling. However, as our findings revealed a reduction in M4-function in response to DA loss, the imbalance of D1-receptor and M4-receptor signaling may contribute to the progression to LID following long-term L-DOPA treatment (Figure 6A). If this is the case, restoring M4-function in dMSNs may be expected to reduce the severity and limit the development of LID (Figure 6A). To test this, we examined the effect of selectively ablating RGS4 from dMSNs in high-dose 6-OHDA lesioned mice treated for 7-8 days with daily i.p. injections of L-DOPA (2 mg/kg) plus benserazide (12 mg/kg) (Figure 6B), a treatment paradigm which elicits therapeutic antiparkinsonian effects (Figure 6E), but also reliable abnormal involuntary movements (AIMs) typical of LID (Figure 6F) in PD mouse models (Shen et al., 2015; Smith et al., 2012).

**Figure 6:**
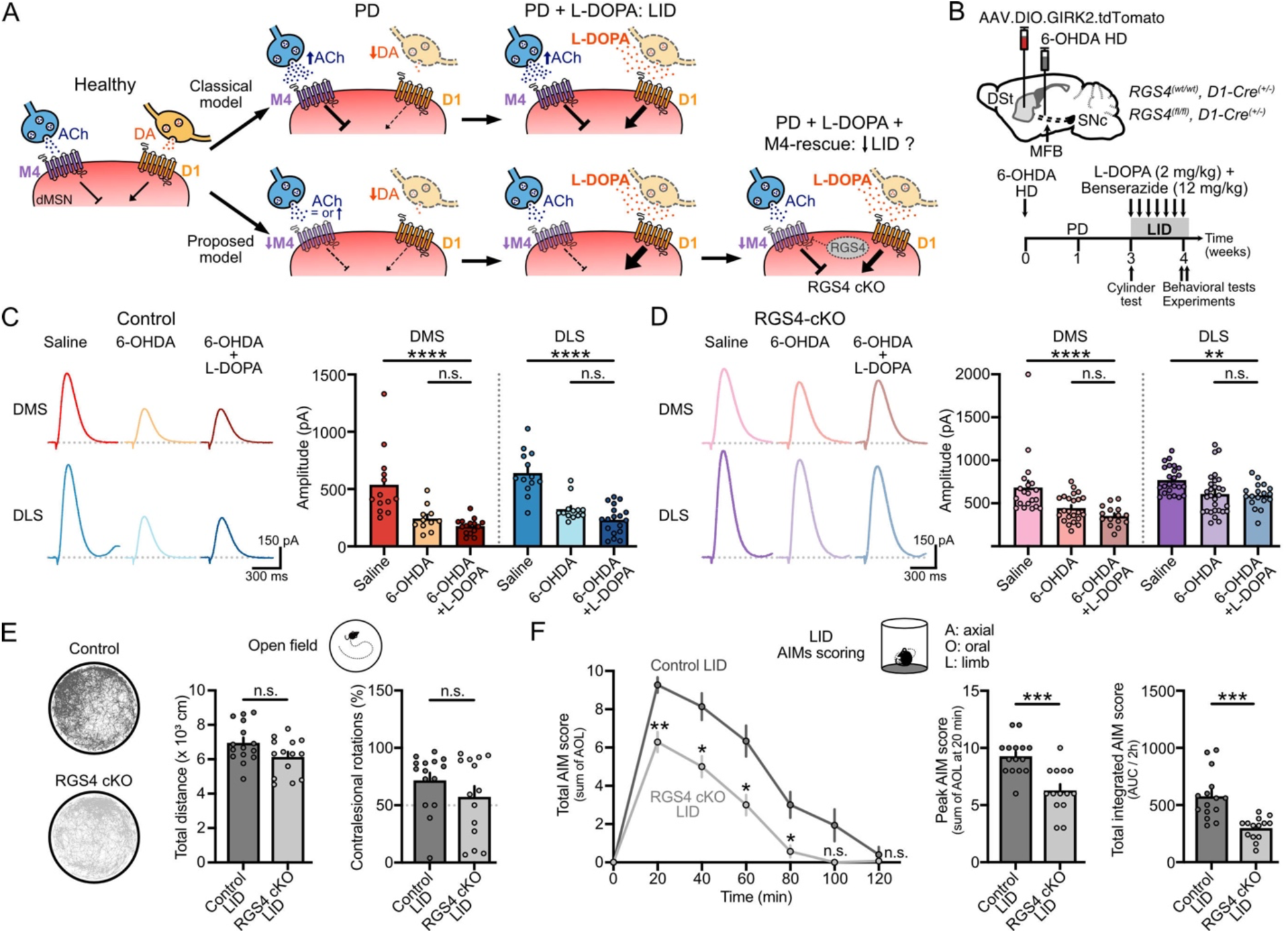
Restoration of M4-transmission alleviates L-DOPA-induced dyskinetic behavior. (A) Cartoon schematics showing the expected changes in the opposite regulation exerted by D1 and M4 receptor signaling pathways in dMSNs in PD and LID according to the classical model (top) and our proposed model (bottom), that set the rationale to examine LID when M4-transmission is restored by knocking-out RGS4. (B) Schematics of AAV9.hSyn.DIO.tdTomato.T2A.GIRK2 and 6-OHDA HD injections into the DSt and MFB respectively in control (RGS4^wt/wt^, D1-Cre^+/-^) and RGS4 cKO in dMSNs mice (RGS4^fl/fl^, D1-Cre^+/-^) (top). Timeline for LID model development and subsequent physiology and behavioral experiments (bottom). (C) Representative traces of electrically evoked M4-IPSCs (25 µA, 0.5 ms) in DMS (top) and DLS (bottom) for saline, 6-OHDA and 6-OHDA+L-DOPA conditions in control mice (left). Amplitude quantification is shown on the bar chart (right). Data from saline and 6-OHDA conditions was taken from Figure 5C (DMS: saline: n=13, N=6; 6-OHDA: n=11, N=6; 6-OHDA+L-DOPA: n=15, N=4 / DLS: saline: n=13, N=7; 6-OHDA: n=13, N=6; 6-OHDA+L-DOPA: n=18, N=5; Kruskal-Wallis, Dunn’s post-hoc test). (D) Representative traces of electrically evoked M4-IPSCs (25 µA, 0.5 ms) in DMS (top) and DLS (bottom) for saline, 6-OHDA and 6-OHDA+L-DOPA conditions in RGS4 cKO mice (left). Amplitude quantification is shown on the bar chart (right). Data from saline and 6-OHDA conditions was taken from Figure 5C (DMS: saline: n=20, N=11; 6-OHDA: n=23, N=11; 6-OHDA+L-DOPA: n=15, N=4 / DLS: saline: n=24, N=11; 6-OHDA: n=29, N=10; 6-OHDA+L-DOPA: n=18, N=4; Kruskal-Wallis, Dunn’s post-hoc test). (E) Example traces of locomotor activity and quantification of total distance (left) and percentage of contralesional rotations in the open field 30 minutes after last L-DOPA administration (control LID: N=15; RGS4 cKO LID: N=14; unpaired t-test for total distance; Mann-Whitney for contralesional rotations). (F) Plot showing total AIM score as a function of time (left) (control LID: N=15; RGS4 cKO LID: N=14; Two-Way RM ANOVA, Šídák’s post-hoc test). Bar graphs show the summary data for the peak of AIM score at 20 min (center) and the integrated AIM score for the total dyskinetic period (2 h) (right) (unpaired t-tests). Summary data is mean ± SEM. n: number of cells, N: number of mice; n.s. p>0.05; *p<0.05; **p<0.01; ***p<0.001; ****p<0.0001. See also Table S1.

We initially began by examining control mice, to see if the 6-OHDA-induced reduction in M4-function could be rescued by prolonged L-DOPA administration. The amplitude of M4-IPSCs from GIRK2^+^ dMSNs from control 6-OHDA lesioned mice (*RGS4^wt/wt^; D1-Cre^+/-^*) treated with L-DOPA, however, was still significantly reduced compared to saline animals and no different than parkinsonian mice not treated with L-DOPA (Figure 6C). This suggests that once initiated, the decrease in M4-function becomes sustained and resistant to changes that replace striatal DA. Similarly, repeating this experiment in RGS4 cKO mice (*RGS4^fl/fl^; D1-Cre^+/-^*) showed that while removal of RGS4 from dMSNs prevented a loss of M4-function following DA lesions, no further potentiation was seen following L-DOPA treatment (Figure 6D). Thus, while the decrease in M4-function can be driven by parkinsonian DA loss, exogenously raising DA levels post-lesion with L-DOPA was unable to revert the aberrant cholinergic transmission.

We next took advantage of the fact that L-DOPA failed to alter M4-mediated cholinergic transmission to evaluate the impact of restoring M4-function on the antiparkinsonian and dyskinetic effects induced by L-DOPA, again hypothesizing that restoring direct pathway M4-function would be expected to limit the severity and development of LID, by balancing the enhanced D1-signaling (Figure 6A). The antiparkinsonian or therapeutic effects of L-DOPA administration were assessed by analyzing locomotor activity and frequency of rotations in an open field arena during a 30 min-observation period in the ‘on’ phase of L-DOPA treatment (Figure 6E). Knockout of RGS4 did not prevent L-DOPA from increasing total distance traveled, velocity or contralesional turning, the latter of which was restored to balanced levels expected for healthy animals (Figures 6E and S5C, and Table S1). Thus, restoration of M4-mediated cholinergic transmission in mice lacking RGS4 did not compromise the therapeutic antiparkinsonian effects of L-DOPA treatment, including enhanced locomotion and rotations.

To directly quantify L-DOPA induced dyskinetic behavior, AIMs for each group were determined by measuring abnormal axial, oral and limb (AOL) movements (Cenci and Lundblad, 2007) (Figure 6F). While L-DOPA-treated control mice rapidly developed dyskinesia, with a total dyskinetic event duration of 120 min, RGS4-lacking mice progressed to dyskinesia at slower rate, terminating the dyskinetic period faster, and with a significant attenuation of the AIMs peak (Figure 6F). Moreover, the overall total AIM score was significantly reduced across the testing period as can be seen by comparing the integrated AIM score or area under the curve for the total dyskinetic period (Figure 6F). Overall, these results indicate that restoring the reduced M4-mediated cholinergic transmission in dMSNs that follows DA loss, ameliorates the dyskinetic behavior in LID without compromising the antiparkinsonian effects of L-DOPA treatment.

## DISCUSSION

According to the DA-ACh balance hypothesis, loss of DA in PD is accompanied by an increase in striatal ACh (Aosaki et al., 2010; Barbeau, 1962; McGeer et al., 1961). The alteration in these two neuromodulators is thought to translate into reduced D1-signaling and enhanced M4-signaling in dMSNs, resulting in an overall inhibition of the direct pathway and suppression of movement. While support for the role of altered cholinergic signaling in PD has come from early ACh dialysis studies (DeBoer et al., 1993) and clinical observations where broad spectrum antimuscarinic drugs partially ameliorate some PD motor symptoms (Betz et al., 2007; Fox et al., 2018; Katzenschlager et al., 2002), directly examining changes in cholinergic transmission in dMSNs following DA depletion has remained unexplored. Here, we examined this issue and found that, as opposed to the classical hypothesis, direct-pathway M4-mediated cholinergic transmission is reduced in response to DA loss in parkinsonian mice. This decrease was the result of impaired postsynaptic M4-function and likely involved both downregulation of receptors and weakened downstream signaling, since either overexpressing M4 receptors or enhancing signaling by selective knockout of RGS4 was sufficient to restore transmission. Restoring impaired M4-transmission improved parkinsonian balance and coordination motor deficits and alleviated LID dyskinetic behaviors. Thus, reduced M4-mediated cholinergic signaling in dMSNs may be a previously unrealized adaptive or compensatory mechanism that contributes to PD symptomatology and LID progression.

Although alterations in cholinergic transmission following DA loss have been predicted, differing results for changes in ACh levels and ChI activity/excitability have been reported, with increases (DeBoer et al., 1993; Ding et al., 2006; Lehmann and Langer, 1983; MacKenzie et al., 1989; Maurice et al., 2015; Sanchez et al., 2011; Sanchez-Catasus et al., 2022; Spehlmann and Stahl, 1976), decreases (Choi et al., 2020; Herrera-Marschitz et al., 1994; McKinley et al., 2019) or lack of changes (Aosaki et al., 1994; Herrera-Marschitz et al., 1990) having been found. When ACh has been directly measured, both increases in extracellular levels with dialysis (DeBoer et al., 1993), and decreases in total striatal content (McKinley et al., 2019) have been reported following induced DA degeneration. Taking advantage of the higher specificity, sensitivity, and spatiotemporal resolution provided by newly developed optical ACh sensors (Jing et al., 2020), we found that while there was a slight increase in the DMS following complete DA lesion, electrically evoked ACh release surprisingly remained unchanged in the DLS, despite this region’s heightened vulnerability to DA loss. Thus, as opposed to more global measures of ACh tone determined using dialysis (DeBoer et al., 1993), the synaptic release of ACh in DLS appears to be unaffected in parkinsonian mice. While other factors, such as alterations in excitability and synaptic inputs may contribute to overall increases in striatal ACh tone when measured *in vivo*, more recent work has found only modest changes in DLS ACh fluctuations following DA lesion in awake behaving animals (Krok et al., 2023). Thus, the hypercholinergic nature of PD remains incompletely understood, and further investigation will be required to determine the alterations that potentially exist in ACh release or tone.

We found that the decrease in direct pathway cholinergic transmission was the result of reduced M4 receptor expression and downstream signaling. This decrease may be an adaptive or compensatory mechanism aimed to counteract the loss of dopaminergic D1-receptor mediated excitatory drive in dMSNs that arises following degeneration of striatal DA inputs. The decrease in M4-expression at the protein level supports past work showing decreased striatal M4 mRNA levels (Kayadjanian et al., 1999; Shan et al., 2015) and M2/M4 radioligand binding (Mann et al., 2018) in PD models (yet see (Laverne et al., 2022; McOmish et al., 2017). The reduced expression of M4 receptors may not be solely explained by structural changes in dendritic arborization or spine density, because these changes are more prominent in iMSNs (Day et al., 2006; Graves and Surmeier, 2019). Instead, as M4 receptor surface levels are reduced in MSNs following prolonged agonist stimulation and by changes in cAMP, the decrease in M4-expression may be also partially driven by altered ACh or DA tone (Bernard et al., 1999; Habecker and Nathanson, 1992; Liste et al., 2002). Besides reduced M4-expression, additional impairments in downstream signaling cascades are also likely involved. We found that RGS4 regulates M4-signaling in dMSNs, and that its ablation is sufficient to restore the decreased M4-transmission following DA loss. As RGS4 transcription appears to be bidirectionally regulated by DA receptor-mediated cAMP signaling, being upregulated when cAMP levels are low and unchanged or downregulated when cAMP increases (Geurts et al., 2002; Pepperl et al., 1998; Taymans et al., 2003), RGS4 mRNA levels can be dynamically and differentially altered in response to acute or prolonged DA depletion (Ding et al., 2006; Geurts et al., 2003; Taymans et al., 2004). Since M4-expression and RGS4 mRNA levels may be differentially regulated across cell types in the striatum (Bernard et al., 1999; Ding et al., 2006), future work is needed to examine specific alterations that may underlie reduced M4-function in dMSNs following DA loss.

As M4 receptor is Gα_i/o_-coupled, its activation by ACh is believed to lead to an inhibition of direct pathway output and a suppression of movement (Foster et al., 2016; Jeon et al., 2010; Moehle et al., 2017; Moehle and Conn, 2019). In partial agreement with this, we found that increasing M4-expression or signaling in dMSNs led to an impaired basal performance in some, but not all, motor tasks in control mice. The surprise from our work is that following DA depletion, increasing direct pathway M4-transmission back to normal levels led to a partial rescue rather than a worsening of motor behavior in parkinsonian mice, supporting a positive role for ACh signaling in motor control. Thus, as opposed to its previously assumed exclusive anti-kinetic function, striatal cholinergic transmission may cooperate to regulate aspects of movement, as has been previously suggested (Avila et al., 2020; Howe et al., 2019). In fact, the alleviation of balance and coordination motor deficits that we observed following rescue of M4-signaling could be related to some features of PD that are refractory or less responsive to DA-replacement therapy (McKay et al., 2019; Sethi, 2008; Wenger et al., 2022), which have been linked to cholinergic system dysfunctions in the striatum and downstream basal ganglia circuits (Bohnen et al., 2022; Pasquini et al., 2021; Wenger et al., 2022). Thus, while our electrophysiology data examined only somatodendritic M4-transmission in striatal dMSNs, the positive behavioral effects resulting from the rescue of M4-signaling may also be partially due to enhanced M4-receptor expression/signaling in axon terminals of dMSNs that project to downstream basal ganglia regions such as the substantia nigra pars reticulata (Moehle et al., 2017).

The need for non-dopaminergic PD therapies lies with treating not only DA-replacement resistant symptoms, but also side effects such as LID that are driven by DA-replacement therapy. As M4 receptors directly oppose D1-signaling in dMSNs (Moehle et al., 2017; Nair et al., 2019; Onali and Olianas, 2002; Shen et al., 2015), we found that restoring M4-signaling back to normal levels by selective ablation of RGS4 alleviated LID abnormal dyskinetic behavior, without compromising the therapeutic antiparkinsonian effects of L-DOPA treatment. This suggests that reductions in M4-mediated transmission may be a previously unrealized factor that accelerates the progression to LID, since loss of M4-signaling in dMSNs may fail to counteract the enhanced D1-receptor signaling that arises with L-DOPA treatment, driving overactivation of the direct pathway and aberrant synaptic depotentiation and plasticity (Feyder et al., 2011; Gerfen, 2003; Girasole et al., 2018; Parker et al., 2018; Picconi et al., 2003, 2003; Ryan et al., 2018; Shen et al., 2015). Because we found similar effects of RGS4 conditional knockout and M4-overexpression on both the rescue of cholinergic transmission and parkinsonian motor impairments, we expect that LID would also be ameliorated by restoring M4 receptor expression. Since RGS4 has a preference for Gα_i/o_-coupled receptors (Berman et al., 1996; Masuho et al., 2020) the effect of knockout likely results from alterations in inhibitory GPCRs such as M4 receptor, however the possibility does exist that, following knockout, alterations in signaling by other GPCRs such at Gα_q_- coupled mGluR5 might also occur (Shen et al., 2015).

While muscarinic antagonists have been used in PD, their utility has been restricted due to side effects from the broad distribution of receptors in the central nervous system and periphery. Despite this, they can be used to treat tremor (Betz et al., 2007) but have little effect on improving bradykinesia and rigidity (Fox et al., 2018; Katzenschlager et al., 2002). Their limited efficacy on treating these core motor symptoms may be explained by the fact that they aim to decrease M4-transmission, which we found is already reduced following DA loss. Therefore, these antagonists may be further inhibiting a signaling pathway that is already downregulated in PD. Muscarinic antagonists have also been used in PD patients undergoing L-DOPA treatment. However their effects on LID are unclear, as there are conflicting results in preclinical models (Bordia et al., 2016; Brugnoli et al., 2020; Chambers et al., 2019; Ding et al., 2011) and in PD patients (Birket-Smith, 1975), varying also according to the ‘on’/’off’ L-DOPA states (Paz and Murer, 2021). Again, since we found that enhancing M4-transmission alleviated dyskinesia, selective agonists rather than antagonists may be preferred as an adjunct therapy to be used with L-DOPA. In line with this, positive allosteric modulators (Brugnoli et al., 2020; Shen et al., 2015), or targeting downstream signaling by RGS4 inhibition (Blazer et al., 2015; Ko et al., 2014), have shown promising results in preclinical models of PD and LID. These same approaches may also be beneficial for improving the balance and coordination deficits in PD that are often associated with L-DOPA-treatment resistant symptoms.

In conclusion, our findings reveal an unexpected reduction in striatal M4-mediated cholinergic transmission in a parkinsonian model, which may constitute a previously unnoticed factor contributing to PD symptomatology and LID progression after prolonged L-DOPA treatment. These results have direct implications for therapeutic development, suggesting that M4-function restoration strategies, as the ones we described here, may be useful as an adjunct to L-DOPA and perhaps other DA-replacement medications.

## Acknowledgments

This work was funded by NIH grants R01-NS95809 (CPF), R01-DA35821 (CPF), Parkinson’s Foundation Postdoctoral Fellowship for Basic Scientists PF-PRF-839073 (BEN), as well as funded in part by Aligning Science Across Parkinson’s (ASAP-020529) (CPF) through the Michael J. Fox Foundation for Parkinson’s Research (MJFF). For the purpose of open access, the author has applied a CC BY public copyright license to all Author Accepted Manuscripts arising from this submission. We thank Drs. Ying Zhu and Jonathan Javitch for providing the AAV.M4-receptor plasmid, Dr. Venetia Zachariou for providing the RGS4-floxed mice, and the University of Colorado Anschutz Behavioral Core.

## Author Contributions

BEN and CPF designed experiments. BEN performed all experiments and data analysis experiments. BEN and CPF wrote the manuscript.

## Data Availability

Datasets are available at: https://doi.org/10.5281/zenodo.8277521 Extended protocols are available at: <u>dx.doi.org/10.17504/protocols.io.dm6gp336jvzp/v1</u>

## MATERIALS AND METHODS

### Experimental Model and Subject Details

#### Animals

All animal experiments were performed in agreement with the protocols approved by Institutional Animal Care and Use Committee (IACUC) at the University of Colorado School of Medicine. Animals were group-housed in a temperature- and humidity-controlled environment on a 12 h light-dark cycle, with water and food available *ad libitum*, and experiments were conducted during the light phase. Both adult male and female 10–11-week-old mice were used for all experiments, including: wildtype C57BL/6J mice (Jackson Laboratory, RRID:IMSR_JAX:000664), Drd1-Cre heterozygote mice (MMRRC, RRID:MMRRC_034258-UCD), and RGS4 conditional knockout (RGS4^fl/fl^:: Drd1-Cre^+/WT^; RGS4 cKO) in D1-MSNs and littermate controls (RGS4^WT/WT^:: Drd1-Cre^+/WT^), which were generated by crossing Drd1-Cre mice with RGS4^fl/fl^ mice provided by Dr. Venetia Zachariou (Avrampou et al., 2019; Han et al., 2010; Stratinaki et al., 2013).

#### Stereotaxic surgery

Mice (7-8 weeks old) were anesthetized with isoflurane, mounted in a stereotaxic frame (Kopf Instruments) and kept under constant 2% isoflurane anesthesia. AAV viruses were unilaterally injected using a Nanoject III (Drummond Scientific) at 2 nL/s into the center of the dorsal striatum, at the following coordinates relative to bregma (in mm): AP +0.9, ML ± 1.85, DV -2.9. Only for experiments aimed to verify regional protein expression levels, injections were conducted separately into the dorsomedial striatum (DMS): AP +0.9, ML ± 1.35, DV -2.9; or the dorsolateral striatum (DLS): AP +0.9, ML ± 2.25, DV -2.9. The needle was kept in the target site for 5 min to allow diffusion. For experiments involving only cell-type specific GIRK expression, 400 nL of AAV9.hSyn.DIO.tdTomato.T2A.GIRK2 (University of Pennsylvania Viral Core, V5688R) were injected. For experiments where GIRK was co-expressed with mAChR M4 or eYFP, 300 nL of AAV9.hSyn.DIO.tdTomato.T2A.GIRK2 were co-injected with 300 nL of AAV9-EF1a-DIO-mChrM4-P2A-eGFP-WPRE-SV40pA (Virovek, provided by Drs. Ying Zhu and Jonathan Javitch) or AAV5.EF1a.DIO.EYFP (Addgene, 27056, RRID:Addgene_27056) respectively. For experiments requiring GRAB_ACh 3.0_, expression, WT mice were injected with 400 nL of AAV9-hSyn-ACh4.3 (ACh3.0) (WZ Biosciences, YL001003-AV9-PUB, RRID:Addgene_121922).

#### Unilateral 6-OHDA mouse model of PD

6-OHDA (6-hydroxydopamine hydrobromide, Sigma-Aldrich, 162957) dissolved in sterile saline was injected (1 µL) unilaterally either at a low dose (1 µg/µL, 6-OHDA LD) or a high dose (4 µg/µL, 6-OHDA HD) into the medial forebrain bundle (MFB) during stereotaxic surgery at the following coordinates relative to bregma (in mm): AP -1.2, ML +1.3, DV -4.75. Mice were pretreated with 25 mg/kg desipramine and 5 mg/kg pargyline dissolved in sterile saline 30 min prior to 6-OHDA injections. To minimize mortality, special care was conducted during at least one-week post-surgery. Briefly, mice remained in a heat pad and received daily sterile saline i.p. injections, food pellets soaked in water and nutritionally fortified water gel (DietGel Recovery, ClearH20).

#### Levodopa-induced dyskinesia mouse model

Three weeks after 6-OHDA injections, mice were examined by the cylinder test to verify DA depletion (See Motor Behavioral Assessment) and then received daily i.p. injections of 2 mg/kg of L-DOPA (3,4-dihydroxy-L-phenylalanine) (Sigma-Aldrich, D9628) plus 12 mg/kg of benserazide hydrochloride (Sigma-Aldrich, B7283) dissolved in sterile saline for 7-8 days. As the extent of dopamine depletion, L-DOPA dose, and treatment duration are factors that aggravate levodopa-induced dyskinesia, which once established is difficult to reverse (Nutt, 2008; Thanvi et al., 2007), a moderate dose and duration of L-DOPA administration were used in an attempt to mimic initial stages of dyskinesia, where potential therapeutic effects could still be tested. Dyskinetic movement was formally scored at the end of the treatment (See Motor Behavioral Assessment). Brain slices were prepared 30min – 1 h after the last L-DOPA administration, when mice were still in the dyskinetic period.

### Method detail

#### Slice preparation

Mice were anesthetized with isoflurane and transcardially perfused with ice-cold sucrose cutting solution containing (in mM): 75 NaCl, 2.5 KCl, 6 MgCl_2_, 0.1 CaCl_2_, 1.2 NaH_2_PO_4_, 25 NaHCO_3_, 2.5 D-glucose and 50 sucrose, bubbled with 95% O_2_ and 5% CO_2_. Coronal striatal slices (240 µm) were cut in the same ice-cold sucrose cutting solution. Slices were then incubated for at least 45 min at 32°C in artificial cerebro-spinal fluid (aCSF) containing (in mM): 126 NaCl, 2.5 KCl, 1.2 MgCl_2_, 2.5 CaCl_2_, 1.2 NaH_2_PO_4_, 21.4 NaHCO_3_, and 11.1 D-glucose, bubbled with 95% O_2_ and 5% CO_2_. 10 µM MK-801 was added to prevent excitotoxicity. After incubation, slices were transferred into a recording chamber and constantly perfused with aCSF warmed to 32 ± 2°C at a rate of 2 mL/min. Neurons were visualized using a BX51WI microscope (Olympus) with an infrared LED, while green and blue LEDs were used for visualizing fluorescence (Thorlabs).

#### Electrophysiology

All recordings were made in the dorsal striatum using Axopatch 200B amplifiers (Molecular Devices). Patch pipettes (1.5 – 2 MΩ) (World Precision Instruments) were made using a pipette puller (Narishige, PC-10). Pipettes for voltage-clamp whole-cell recordings from MSNs contained (in mM): 135 D-Gluconate (K), 10 HEPES(K), 0.1 CaCl_2_, 2 MgCl_2_ and 10 BAPTA-tetra potassium, with 1 mg/mL ATP, 0.1 mg/mL GTP, and 1.5 mg/mL phosphocreatine (pH 7.35, 275 mOsm). MSNs were held at -60 mV. To reduce the variability of GIRK outward currents between cells and animals, all recordings from GIRK^+^ MSNs were conducted in regions showing robust reporter fluorescence (Gong et al., 2021; Zych and Ford, 2022), and where indicated, ACh release was triggered by electrical stimulation using a monopolar glass stimulating electrode filled with aCSF, positioned consistently 200 µm away from the recorded cell (0.5 ms). No series resistance compensation was applied, and cells were discarded if series resistance was ≥15 MΩ. For paired ChI-dMSN recordings, the presynaptic ChI internal solution contained (in mM): 135 D-Gluconate (K), 10 HEPES(K), 0.1 CaCl_2_, 2 MgCl_2_ and 0.1 EGTA, with 1 mg/mL ATP, 0.1 mg/mL GTP, and 1.5 mg/mL phosphocreatine (pH 7.35, 275 mOsm). All putative ChIs were identified by their large size and presence of a hyperpolarization activated inward current (H-current) with a hyperpolarization protocol (30 mV, 5 s). Recordings were acquired with Axograph 1.76 (Axograph Scientific; RRID SCR_014284) at 10 kHz and filtered to 2 kHz, or with LabChart (ADInstruments) at 1 kHz. Unless otherwise noted, all drugs were bath applied and recordings were performed in aCSF containing 10 µM DNQX, 100 µM picrotoxin, 300 nM CGP 55845, 10 mM SCH 23390, 10 µM DHβE and 200 nM sulpiride in order to isolate muscarinic cholinergic transmission.

#### 2-photon imaging GRAB_ACh 3.0_ recordings

2-photon imaging was performed using a 2-photon laser scanning microscopy system, custom-built on a BX51WI microscope (Olympus). A Ti:Sapphire laser (Chameleon Ultra I; Coherent) was tuned to emit pulsed excitation at 920 nm and scanned using a pair of X-Y galvanometer mirrors (6215, Cambridge Technology). Emitted fluorescence was collected through a water-immersion objective (60X, Olympus), a dichroic mirror (T700LPXXR, Chroma) and filters (ET680sp and ET525/50 m-2P, Chroma), and was detected using a GaAsP photomultiplier tube (PMT, H10770PA-40, Hamamatsu). A current preamplifier (SR570, Stanford Research Systems) was used to convert the output to voltage, which was then digitized by a data acquisition card (PCI-6110, National Instruments).

For ACh imaging experiments, fluorescence changes of the GRAB_ACh3.0_ sensor were measured using 2-photon non-raster scanning photometry and a custom software (Toronado; https://github.com/StrowbridgeLab/Toronado-Laser-Scanning) as previously described (Pressler and Strowbridge, 2019). The laser was repeatedly scanned across a small circular path (150 nm diameter) at a selected region of interest, and fluorescence was continuously collected from that spot. The PMT signal was converted by the same preamplifier (SR570, Stanford Research Systems; sensitivity 100 nA/V), but further filtered to 500 Hz with the gain increased two-fold (FLA-01, Cygnus Technologies). Then the signal was acquired using a data acquisition device (ITC-18, HEKA Instruments) and recorded using Axograph X (Axograph Scientific).

ACh release was triggered by electrical stimulation with the electrode positioned in the center of a square region of interest (25 µm x 25 µm), and the average change in GRAB_ACh3.0_ fluorescence was quantified in Fiji (ImageJ, RRID:SCR_002285). For paired-pulse ratio experiments, GRAB_ACh3.0_ fluorescence was collected from individual spots of interest inside the square region during paired electrical stimulation with 100 ms inter stimulus interval (ISI).

To verify equal ACh sensor expression levels across striatal regions, the GRAB_ACh3.0_ fluorescence signal was recorded every 30s in the square region of interest for a 5 min-period of baseline and then following bath application of saturating concentrations of ACh (100 µM) and an AChE inhibitor ambenonium (100 nM).

#### Immunohistochemistry and fluorescence imaging

Mice were anesthetized using isoflurane and perfused transcardially with cold PBS followed by cold 4% paraformaldehyde in PBS, pH 7.4. Brains were post-fixed in 4% PFA at 4°C for additional 2h, equilibrated in 30% sucrose solution for 2 days, and rapidly frozen in embedding freezing medium (Fisher Scientific). Dorsal striatum and/or midbrain coronal slices of 30 µm thickness were obtained using a Leica CM1950 cryostat (Leica Microsystems).

For tyrosine hydroxylase immunohistochemistry, sections were mounted on slides and blocked in 5% normal donkey serum in PBS-T (0.3% Triton X-100) for 1 hour at RT. Slides were then washed in PBS and incubated with rabbit anti-TH primary antibody (1:200, Millipore AB152, RRID: AB_390204) overnight at 4°C. After PBS washes, slides were incubated with donkey anti-rabbit Alexa Fluor 488 secondary antibody (1:500, Life Technologies A21206, RRID: AB_2535792) for 1 hour at RT and washed afterward with PBS.

For M4 receptor immunohistochemistry, sections were mounted on slides and stained using a monoclonal mouse anti-muscarinic acetylcholine receptor M4/CHRM4 primary antibody (1:500, abcam, ab77956, RRID: AB_1566454), with blocking reagents and secondary anti-mouse biotinylated antibody from the M.O.M. Immunodetection kit (Vector Laboratories, BMK-2202, RRID:AB_2336833) according to manufacturer directions. Sections were then incubated with streptavidin Alexa Fluor 594 conjugate (1:1000, Invitrogen, S32356) for 30 min at RT, and washed afterward with PBS. Following immunostaining, sections were finally mounted with an anti-fading mounting media. When only visualizing fluorescence reporters, sections were mounted right after being washed in PBS without additional immunostaining. Fluorescent images were acquired using a slide scanner microscope (VS120, Olympus) and processed in Fiji (ImageJ, RRID:SCR_002285).

#### Western Blot

Dorsal striatum tissue for western blot analysis was isolated as previously described (Gong et al., 2021; Marcott et al., 2014) from saline and 6-OHDA injected hemispheres, and then further dissected with a sagittal section along the midline of the dorsal striatum to separate DMS and DLS regions. The samples were homogenized and denatured with STE buffer (10 mM Tris-Cl pH 7.5, 1 mM EDTA pH 8.0, 1% SDS) at 100°C for 5 min. Protein concentrations were quantified by Pierce BCA Protein Assay (ThermoScientific, 23225-7). Equivalent amounts of protein were subjected to SDS-PAGE on 10% polyacrylamide gels, and then transferred to methanol activated PVDF membranes (Perkin Elmer). Blots were blocked with 5% milk in TBS-T (0.1% Tween) for 1h at RT and then immunolabeled with rabbit primary antibodies, anti-GIRK2 (1:500, Alomone labs, APC-006, RRID: AB_2040115) and anti-actin (1:2000, Cell Signaling Technology, 4970S, RRID: AB_2223172) overnight at 4°C in blocking buffer. Blots were then probed with horseradish peroxidase (HRP)-conjugated secondary antibody (1:3000, GE Healthcare, NA934, RRID: AB_772206) for 1h at RT. Proteins were detected using a chemiluniscent substrate (SuperSignal West Pico PLUS Chemiluniscent Substrate, ThermoScientific, 34577) and visualized in FluorChem SP (Alpha Innotech). Densitometry analysis was performed in Fiji (ImageJ, RRID:SCR_002285).

#### Chemicals

Picrotoxin (1128), (+)-MK-801 maleate (0924), DNQX (0189), DHβE hydrobromide (2349), SCH 23390 hydrochloride (0925), (S)-(-)-sulpiride (0895), acetylcholine chloride (2809), oxotremorine M (1067), VU 0255035 (3727), tropicamide (0909), ambenonium dichloride (0388), and scopolamine hydrobromide (1414) were obtained from Tocris Bioscience. EGTA (E4378), desipramine hydrochloride (D3900), 6-hydroxydopamine hydrobromide (162957), 3,4-dyhidroxy-L-phenylalanine (D9628), benserazide hydrochloride (B7283) were from Sigma-Aldrich. CGP55845 hydrochloride (HB0960) was purchased from Hello Bio, BAPTA tetra potassium salt from Invitrogen (B1204) and pargyline hydrochloride from Abcam (ab141265).

#### Motor behavioral assessment

Prior to each behavioral test, mice were habituated to the testing room for 30-40 min. All behavioral tests were performed 3 weeks following stereotaxic injections. Video recordings were analyzed blindly.

#### Cylinder test

Forelimb use asymmetry during exploratory activity was assessed by cylinder test. Individual mice were placed in a clear plastic cylinder (10.5 cm diameter; 14.5 cm height) with mirrors located behind for appropriate vision and were video recorded for 5 min for later post-hoc scoring. No prior habituation was allowed before video recording and only wall contacts executed with fully extended digits were scored. Data was expressed as a percentage of touches performed with the forelimb contralateral to the injected side with respect to the total paw use (ipsilateral + contralateral).

Forelimb lateralization revealed by this test was conducted to verify DA-depletion prior to further experiments. Only mice whose contralateral paw use was lower than 45% and 40% for 6-OHDA LD and HD conditions respectively were included for experiments.

#### Rotarod

Rotarod test was performed to assess motor balance and coordination, using an accelerated protocol from 3 to 30 rpm in 5 min (Med Associates). Each mouse underwent 3 trials on the same day with a separation of 10 min between trials, without previous training. The latency to fall or time to reach maximum speed was recorded, and the data was expressed as the average of the two best trials.

#### Balance beam

A balance beam test was conducted to assess motor balance and coordination by measuring mice ability to transverse a graded series of beams to reach a goal cage. Animals were trained for one day in the medium size square beam (12 mm), and the following day, that one and two additional beams (19 mm and 6 mm) were presented to the mice. Animals were video recorded while undergoing two trials per beam size, and the number of slips and time to cross the beam were registered, with a maximal score of 60 s (unsuccessful trial). The data was expressed as the average of the two trials for each beam size. Number of slips were only counted for successful trials (<60s).

#### Open field

Locomotion was assessed using the open field test. Each mouse was gently placed into the square arena (50 x 50 cm) and video recorded for 10 min using an overhead camera. Tracking and post-hoc analysis of total distance and velocity were conducted with Ethovision XT 17.5 (Noldus, RRID:SCR_004074).

For a subset of mice used for LID model, open field test was assessed after 7-8 days of daily L-DOPA injections. Each mouse received the last L-DOPA i.p. injection and was placed into a clear plastic cylinder (25.4 cm diameter; 30.5 cm height). 30 min later, mice were video recorded for another 30 min-period using an overhead camera. Tracking and post-hoc analysis of locomotion and rotations (threshold: 90°, minimum distance travelled: 2 cm, (Ryan et al., 2018)) were automatically performed with Ethovision XT 17.5 (Noldus, RRID:SCR_004074).

#### Abnormal Involuntary Movement score

Dyskinesia was assessed using the Abnormal Involuntary Movement score (AIMs) (Cenci and Lundblad, 2007). Mice were individually placed in clear plastic cylinders (20 cm diameter; 25.5 cm height) following L-DOPA injection and abnormal axial (A), orolingual (O) and limb (L) movements were video recoded for 1 min every 20 min for a total length of 2h. The AIM scale ranges from 0 to 4 for each body segment during a 1-minute period: 0-normal movement, 1-abnormal movement for <50% of the time, 2-abnormal movement for >50% of the time, 3-abnormal movement for the entire period that can be interrupted by sensory stimuli, and 4-continuous abnormal movement, uninterruptible. Total AIM score is the sum of scores for AOL, being 12 the maximum score in 1 minute. An integrated AIM score was calculated as the area under the curve (AUC) in a plot of total AIM score vs time for the duration of the dyskinetic episode (2h) (Winkler et al., 2002).

The video recordings were blindly acquired and analyzed post-hoc by the experimenter.

### Quantification and statistical analysis

#### Statistics

Statistical analyses were performed in Prism 10 (GraphPad 10.0.2, RRID:SCR_00306). All data are shown as mean ± SEM.

Data sets that passed the Shapiro-Wilk test for normality were analyzed using parametric tests, otherwise, non-parametric tests were applied. For comparison between two groups, the following two-tailed statistical tests were conducted as appropriate: unpaired or paired Student t test, Mann-Whitney U test or Wilcoxon matched-pairs signed rank test. For comparisons between more than two groups, the following statistical tests were applied as appropriate: One-way analysis of variance (ANOVA), Kruskal-Wallis ANOVA, Two Way ANOVA or Mixed-model ANOVA with Geisser-Greenhouse’s correction. Repeated measures (RM) version for those tests were performed for matched or paired data. When significant differences were found in ANOVA tests, post-hoc multiple comparison tests were performed as appropriate, including Dunnet’s, Dunn’s, Tukey’s, or Holm-Sidak’s tests. Concentration-response curves were adjusted by nonlinear regression (Hill coefficient = 1) and EC_50_ was assumed to be Gaussian distributed. Statistical significance was established as p≥0.05 (n.s.), p<0.05 (*), p<0.01 (**), p<0.001 (***), and p<0.0001 (****). ‘n’ denotes number of cells and ‘N’ represents number of animals. p-values, n, N, and specific statistical tests used for each experiment, as well as for post-hoc multiple comparisons are stated in the figure legends and Supplemental Table S1.

## SUPPLEMENTAL FIGURES LEGENDS

**Figure S1: (associated to Figure 1).**
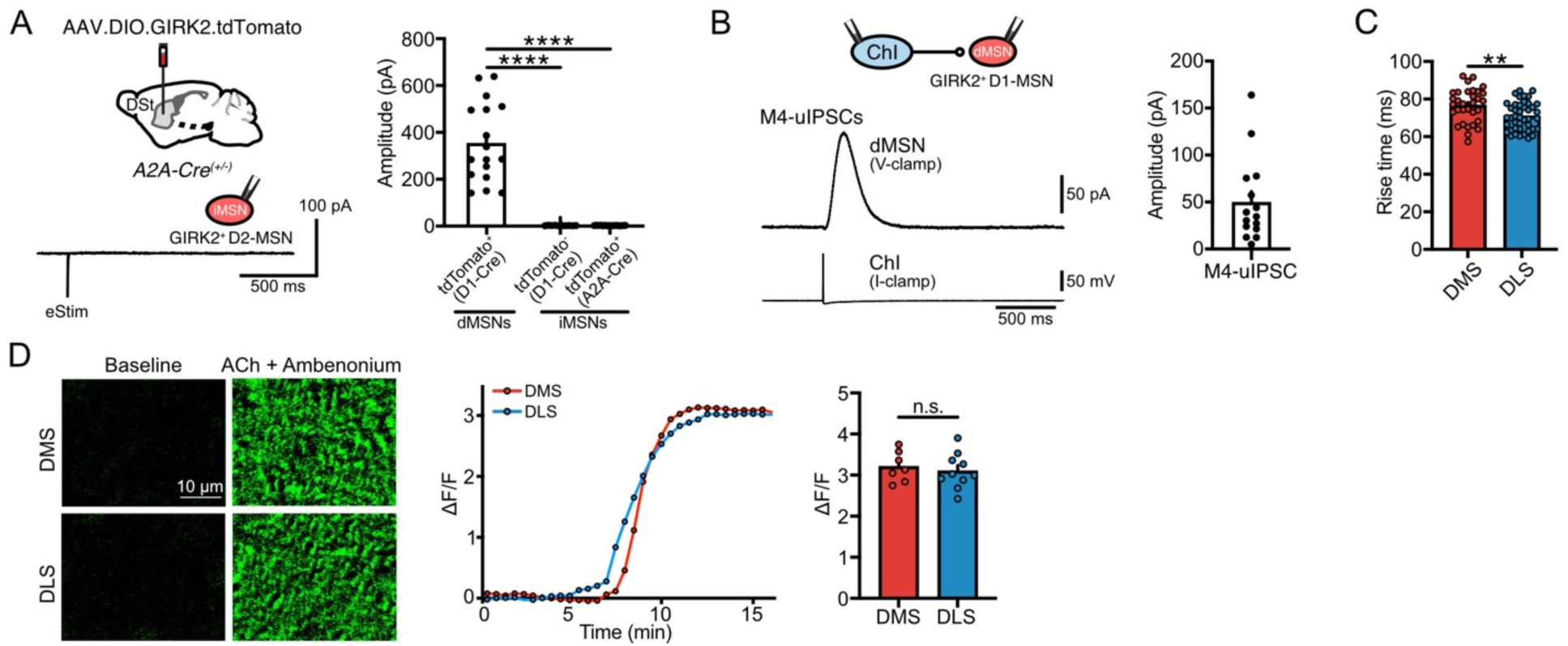
(A) Schematic of AAV9.hSyn.DIO.tdTomato.T2A.GIRK2 injection into the DSt of A2A-Cre mice (top left) and representative recording of tdTomato^+^ iMSNs showing the absence of M4-IPSCs after electrical stimulation (bottom left). Quantification is shown in the bar chart (right). For comparison purposes, first and second bars are control data taken from Figure 1C (tdTomato^+^ dMSNs from D1-Cre mice: n=17, N=6; tdTomato^-^ ’iMSNs’ from D1-Cre mice: n=10, N=3; tdTomato^+^ iMSNs from A2A-Cre mice: n=15, N=3; Kruskal-Wallis; Dunn’s post-hoc test). (B) Representative traces and summary data of ChI-dMSN paired recordings, with ChI and dMSN in current-clamp and voltage-clamp modes respectively. An action potential in ChI elicits a time-locked unitary M4-IPSC in paired dMSN (M4-uIPSCs control: n=15, N=6). (C) Summary data of rise time (from 10 to 90%) across striatal regions (DMS: n=32, N=13; DLS: n=43, N=16; unpaired t-test). (D) Representative fluorescent changes of GRAB_ACh3.0_ in DMS and DLS following bath application of 100 µM ACh and 100 nM ambenonium to control sensor expression levels (left). Plot showing representative time courses of ΔF/F for each striatal region (center). Summary data for maximal ΔF/F (right) (DMS: n=7, N=4; DLS: n=10, N=5; unpaired t-test). Summary data is mean ± SEM. n: number of cells, N: number of mice; n.s. p>0.05; *p<0.05; **p<0.01; ***p<0.001; ****p<0.0001. See also Table S1.

**Figure S2: (associated to Figure 3).**
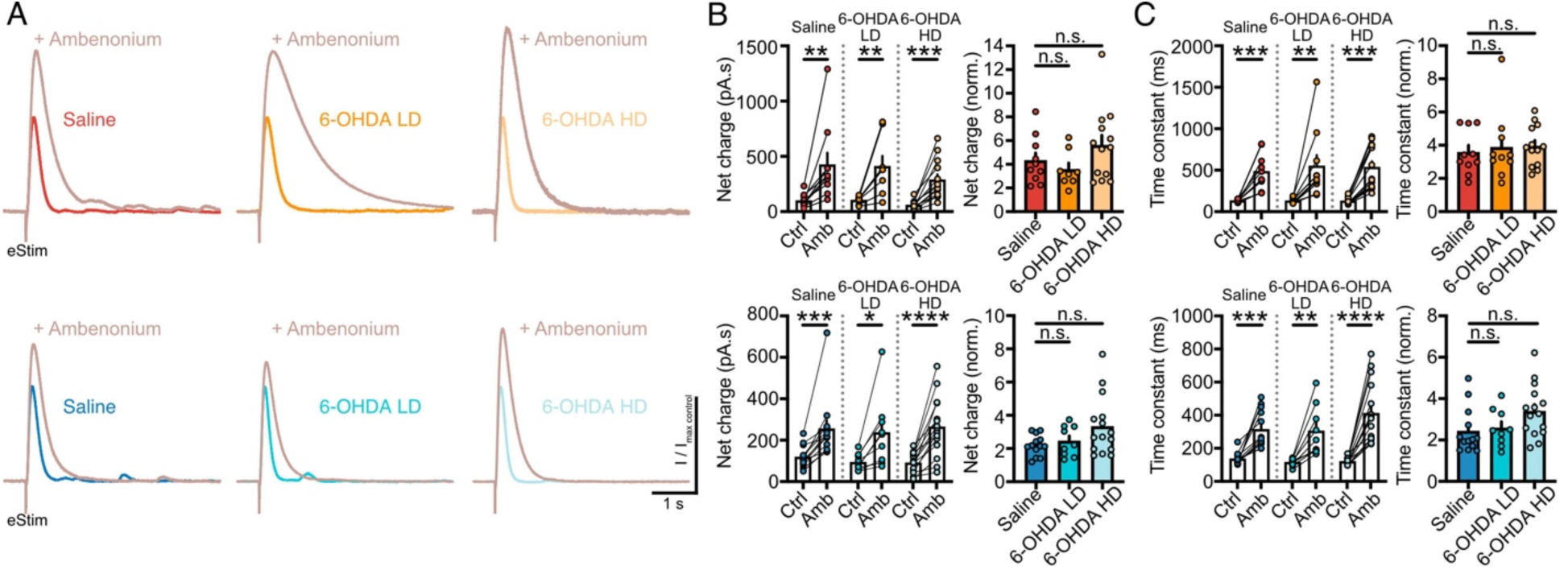
(A) Representative traces of electrically evoked M4-IPSCs (25 µA, 0.5 ms) in DMS (top) and DLS (bottom) in saline, 6-OHDA LD and 6-OHDA HD conditions, before (Ctrl) and after (Amb) bath application of a non-saturating concentration of ambenonium (10 nM). (B) Quantification of net charge, absolute values (left) and normalized to control (right), following ambenonium application for saline, 6-OHDA LD and 6-OHDA HD conditions in DMS (top) (saline n=10, N=6, Wilcoxon; 6-OHDA LD n=8, N=6, paired t-test; 6-OHDA HD n=13, N=8, paired t-test / For normalized data: One-Way ANOVA) and DLS (bottom) (saline n=13, N=7, Wilcoxon; 6-OHDA LD n=9, N=6, paired t-test; 6-OHDA HD n=15, N=8, paired t-test / For normalized data: Kruskal-Wallis). (C) Quantification of time constant, absolute values (left) and normalized to control (right), following ambenonium application for saline, 6-OHDA LD and 6-OHDA HD conditions in DMS (top) (saline n=10, N=6, paired t-test; 6-OHDA LD n=10, N=6, Wilcoxon; 6-OHDA HD n=13, N=8, Wilcoxon / For normalized data: Kruskal-Wallis) and DLS (bottom) (saline n=13 N=7, Wilcoxon; 6-OHDA LD n=9, N=6, paired t-test; 6-OHDA HD n=15, N=8, Wilcoxon / For normalized data: Kruskal-Wallis). Summary data is mean ± SEM. n: number of cells, N: number of mice; n.s. p>0.05; *p<0.05; **p<0.01; ***p<0.001; ****p<0.0001. See also Table S1.

**Figure S3: (associated to Figure 4).**
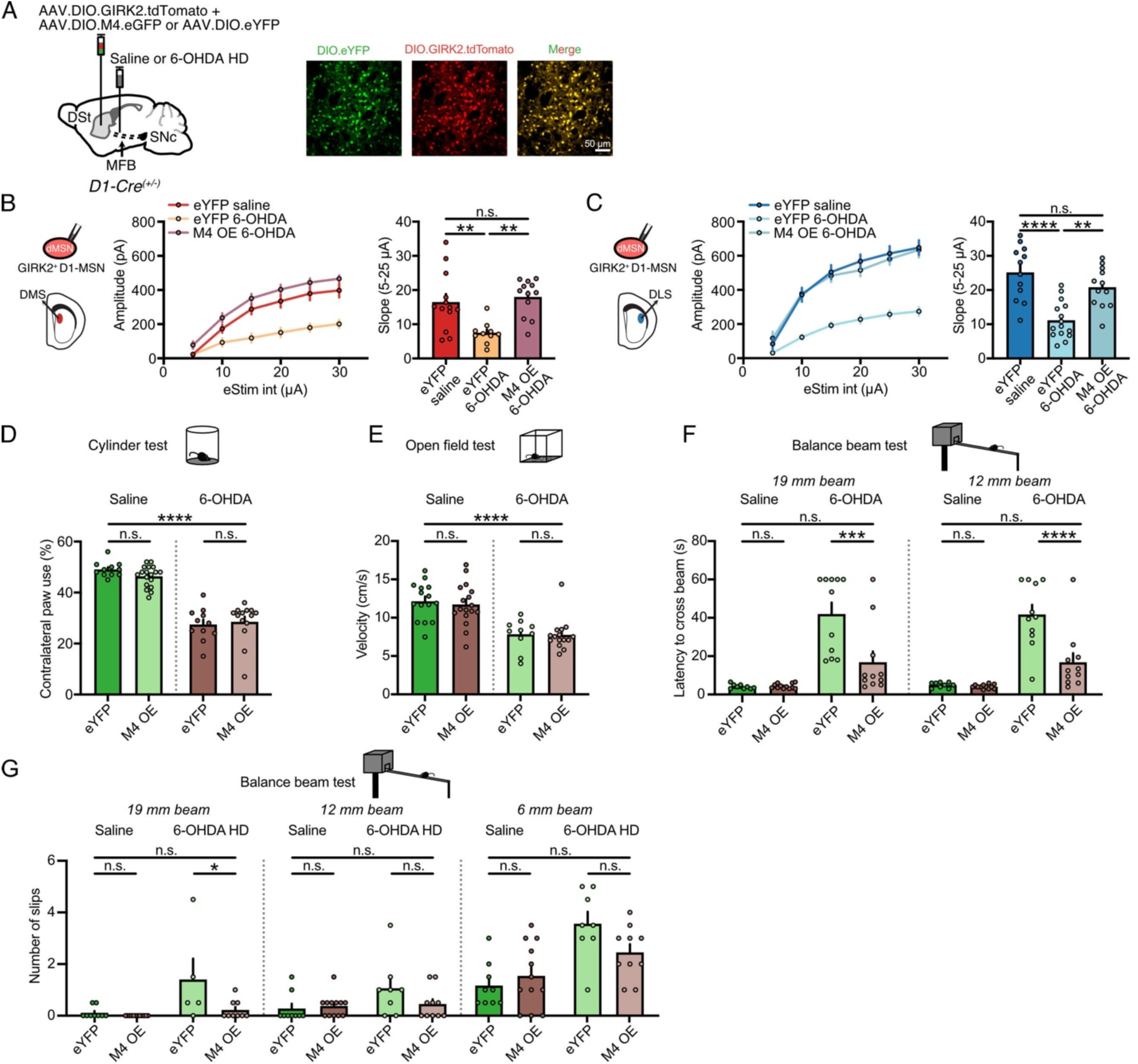
(A) Schematics of AAV.DIO.M4.eGFP or AAV.DIO.eYFP co-injections with AAV9.hSyn.DIO.tdTomato.T2A.GIRK2 into the DSt, and saline/6-OHDA HD injections into the MFB of D1-Cre mice (left). Close-up images of dMSNs co-expressing eYFP and tdTomato fluorescent reporters (right). (B) Plot of M4-IPSCs amplitudes versus electrical stimulation intensity (left) and summary data of slope for 5-25 µA range (right) in DMS for eYFP saline, eYFP 6-OHDA and M4 OE 6-OHDA conditions (eYFP saline: n=13, N=6; eYFP 6-OHDA: n=10, N=5; M4 OE 6-OHDA: n=12, N=6; One-Way ANOVA; Tukey’s post-hoc test). (C) Plot of M4-IPSCs amplitudes versus electrical stimulation intensity (left) and summary data of slope for 5-25 µA range (right) in DLS for eYFP saline, eYFP 6-OHDA and M4 OE 6-OHDA conditions (eYFP saline: n=12, N=6; eYFP 6-OHDA HD: n=15, N=5; M4 OE 6-OHDA HD: n=12, N=7; One-Way ANOVA; Tukey’s post-hoc test). (D) Summary data of cylinder test performance (% contralateral paw use) (eYFP saline: N=11; M4 OE saline: N=18; eYFP 6-OHDA: N=11; M4 OE 6-OHDA: N=14; Two-Way ANOVA, Šídák’s post-hoc test). (E) Summary data of mean velocity in open field test (eYFP saline: N=15; M4 OE saline: N=17; eYFP 6-OHDA: N=10; M4 OE 6-OHDA: N=15, Two-Way ANOVA, Šídák’s post-hoc test). (F) Summary data of latency to cross 19 mm- and 12 mm-beams in balance beam test (eYFP saline: N=9; M4 OE saline: N=12; eYFP 6-OHDA: N=11; M4 OE 6-OHDA: N=11; Two-Way ANOVA, Šídák’s post-hoc test). (G) Average number of foot slips in balance beam test for all beams (eYFP saline: N=9; M4 OE saline: N=12; eYFP 6-OHDA: N=5-8; M4 OE 6-OHDA: N=9-10; Two-Way ANOVA, Šídák’s post-hoc test). Summary data is mean ± SEM. n: number of cells, N: number of mice; n.s. p>0.05; *p<0.05; **p<0.01; ***p<0.001; ****p<0.0001. See also Table S1.

**Figure S4: (associated to Figure 5).**
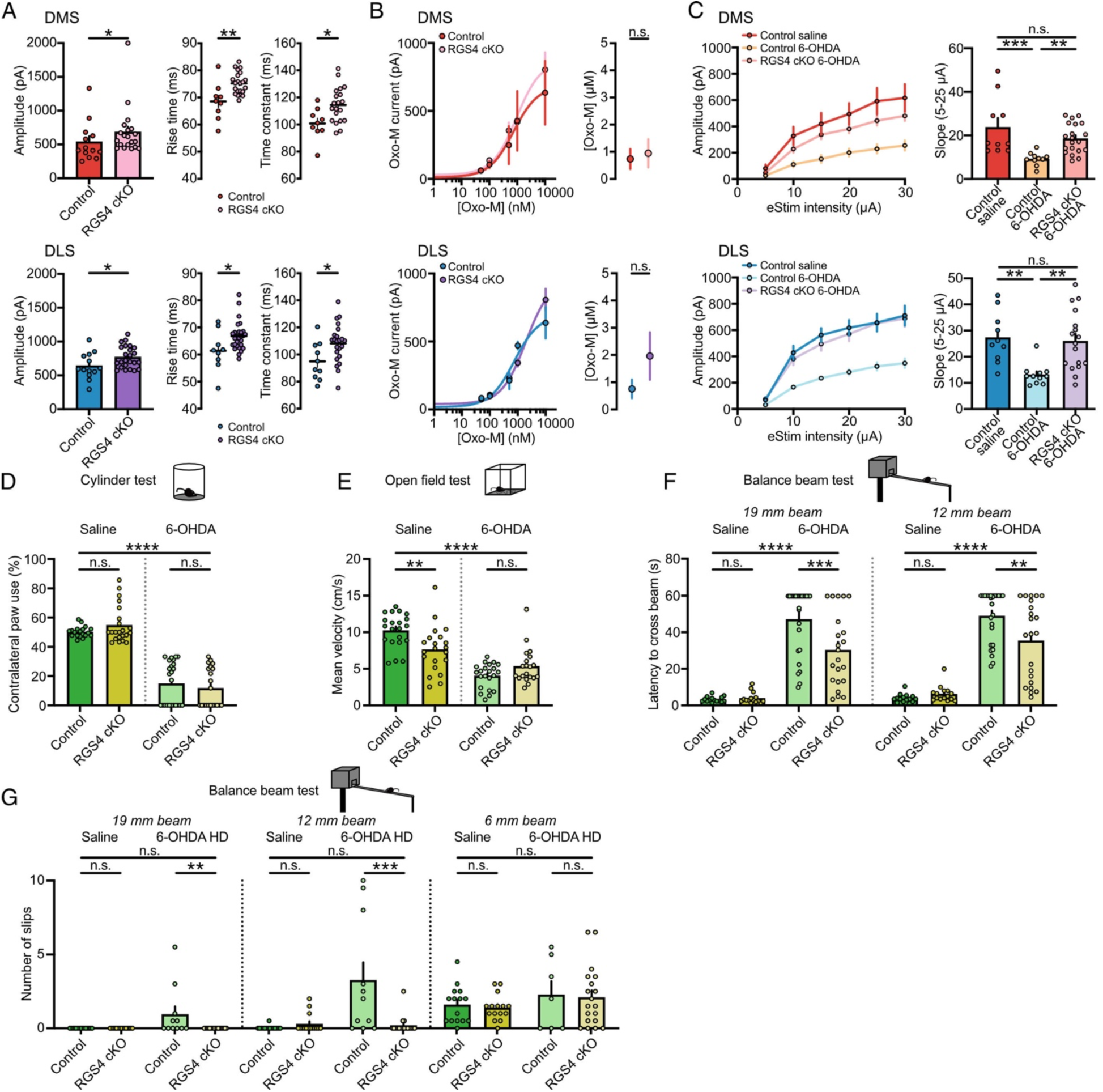
(A) Quantification of electrically evoked M4-IPSCs amplitudes (25 µA, 0.5 ms), rise time and time constant (15 µA, 0.5 ms) in DMS (top) and DLS (bottom) for control and RGS4 cKO saline conditions (Amplitudes (data taken from Figure 5C) - DMS: control saline: n=13, N=6; RGS4 cKO saline: n=20, N=11; Mann-Whitney - DLS: control saline: n=13, N=7; RGS4 cKO saline: n=24, N=11; unpaired t-test / Rise time -DMS: control saline: n=9, N=6; RGS4 cKO saline: n=22, N=11; unpaired t-test -DLS: control saline: n=9, N=5; RGS4 cKO saline: n=28, N=11; Mann-Whitney / Time constant -DMS: control saline: n=9, N=6; RGS4 cKO saline: n=20, N=11; unpaired t-test -DLS: control saline: n=10, N=6; RGS4 cKO saline: n=23, N=11; unpaired t-test). (B) Oxotremorine concentration-response curve for M4 receptors in DMS (top) and DLS (bottom) for control and RGS4 cKO saline conditions (left) with EC_50_ values (middle) (DMS: control saline: n=21, N=4-5; RGS4 cKO saline: n=31, N=4-10; unpaired t-test / DLS: control saline: n=22, N=4-5; RGS4 cKO saline: n=29, N=4-10; unpaired t-test). (C) Plot of M4-IPSCs amplitudes versus electrical stimulation intensity (left) and summary data of slope for 5-25 µA range (right) in DMS (top) and DLS (bottom) for control saline, control 6-OHDA and RGS4 cKO 6-OHDA conditions (DMS: control saline: n=10, N=6; control 6-OHDA: n=10, N=6; RGS4 cKO 6-OHDA: n=20, N=11 / DLS: control saline: n=10, N=6; control 6-OHDA: n=11, N=6; RGS4 cKO 6-OHDA: n=17, N=8; Kruskal-Wallis, Dunn’s post-hoc test). (D) Summary data of cylinder test performance (% contralateral paw use) (control saline: N=19; RGS4 cKO saline: N=22; control 6-OHDA: N=24; RGS4 cKO 6-OHDA: N=21; Two-Way ANOVA, Šídák’s post-hoc test). (E) Summary data of mean velocity in open field test (control saline: N=21; RGS4 cKO saline: N=21; control 6-OHDA: N=22; RGS4 cKO 6-OHDA: N=20; Two-Way ANOVA, Šídák’s post-hoc test). (F) Summary data of latency to cross 19 mm- and 12 mm-beams in balance beam test (control saline: N=15; RGS4 cKO saline: N=15; control 6-OHDA: N=26; RGS4 cKO 6-OHDA: N=22; Two-Way ANOVA, Šídák’s post-hoc test). (G) Average number of foot slips in balance beam test for all beams (control saline: N=15; RGS4 cKO saline: N=15; control 6-OHDA: N=7-11; RGS4 cKO 6-OHDA: N=17-19; Two-Way ANOVA, Šídák’s post-hoc test). Summary data is mean ± SEM. n: number of cells, N: number of mice; n.s. p>0.05; *p<0.05; **p<0.01; ***p<0.001; ****p<0.0001. See also Table S1.

**Figure S5 (associated to Figure 6).**
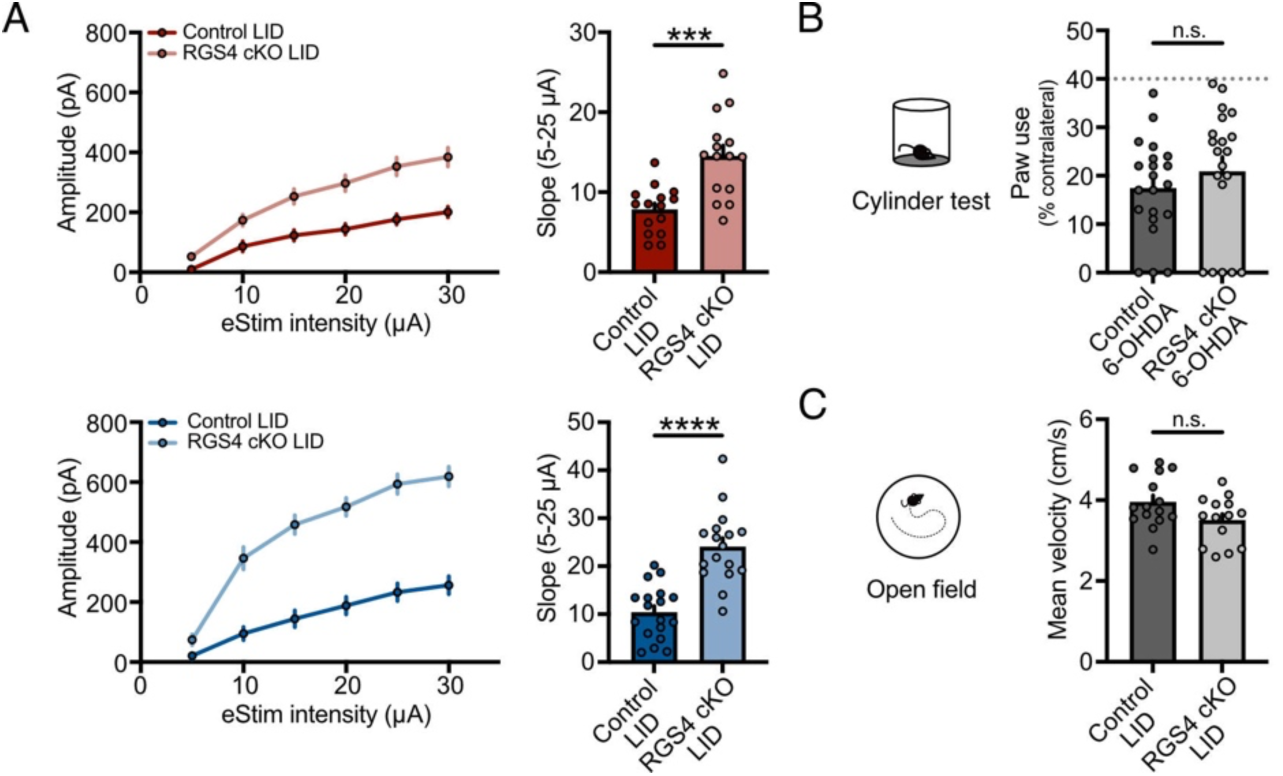
(A) Plot of M4-IPSCs amplitudes versus electrical stimulation intensity (left) and summary data of slope for 5-25 µA range (right) in DMS (top) and DLS (bottom) for control LID and RGS4 cKO LID conditions (DMS: control LID: n=15, N=4; RGS4 cKO LID: n=15, N=4; unpaired t-test / DLS: control LID: n=18, N=5; RGS4 cKO LID: n=17, N=4; unpaired t-test). (B) Summary data of cylinder test performance (% contralateral paw use) for control and RGS4 cKO 6-OHDA HD conditions to assess DA depletion before chronic L-DOPA administration (control 6-OHDA: N=21; RGS4 cKO 6-OHDA: N=20; Mann-Whitney). (C) Quantification of mean velocity in the open field 30 minutes after last L-DOPA administration (control LID: N=15; RGS4 cKO LID: N=14; unpaired t-test). Summary data is mean ± SEM. n: number of cells, N: number of mice; n.s. p>0.05; *p<0.05; **p<0.01; ***p<0.001; ****p<0.0001. See also Table S1.

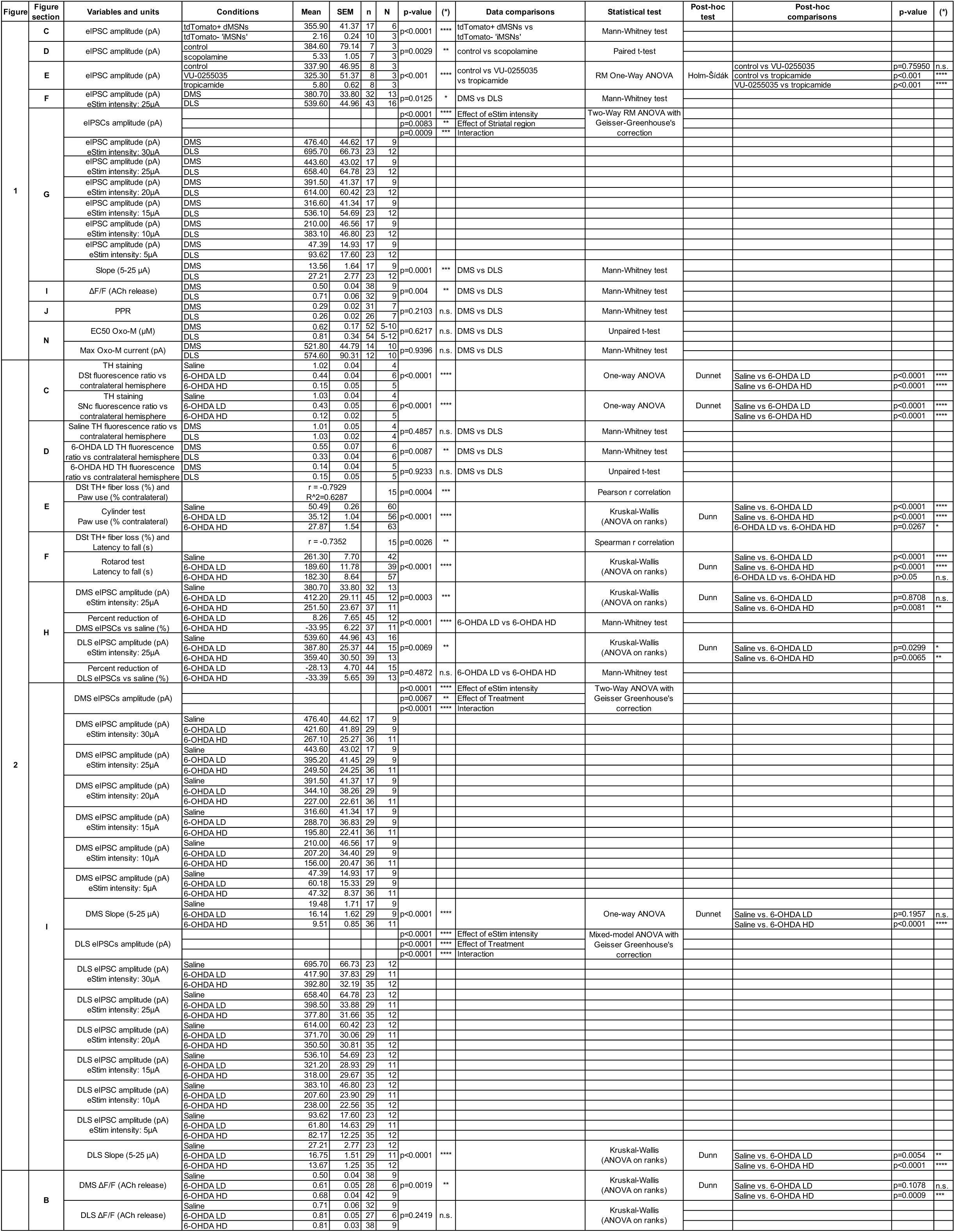

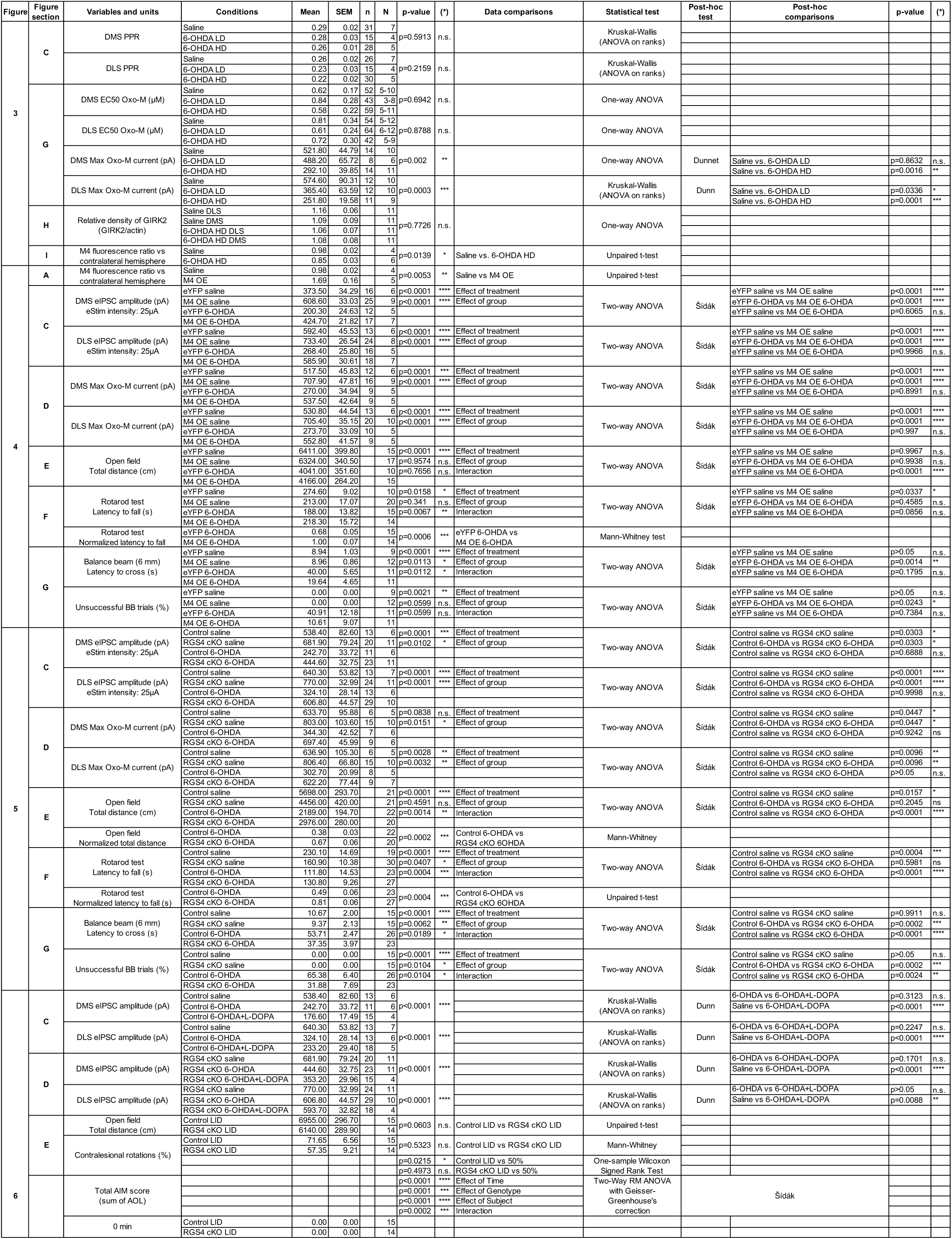

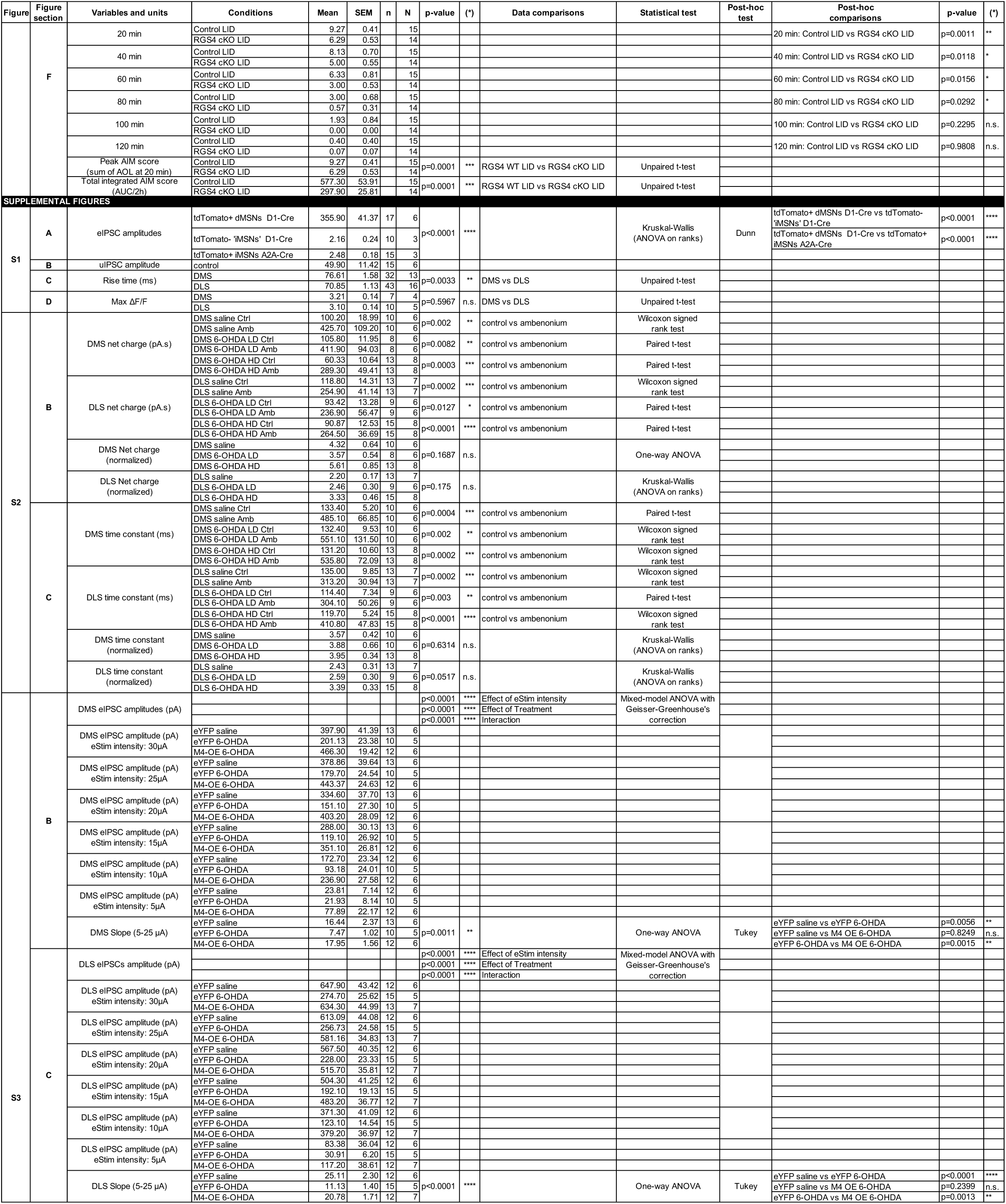

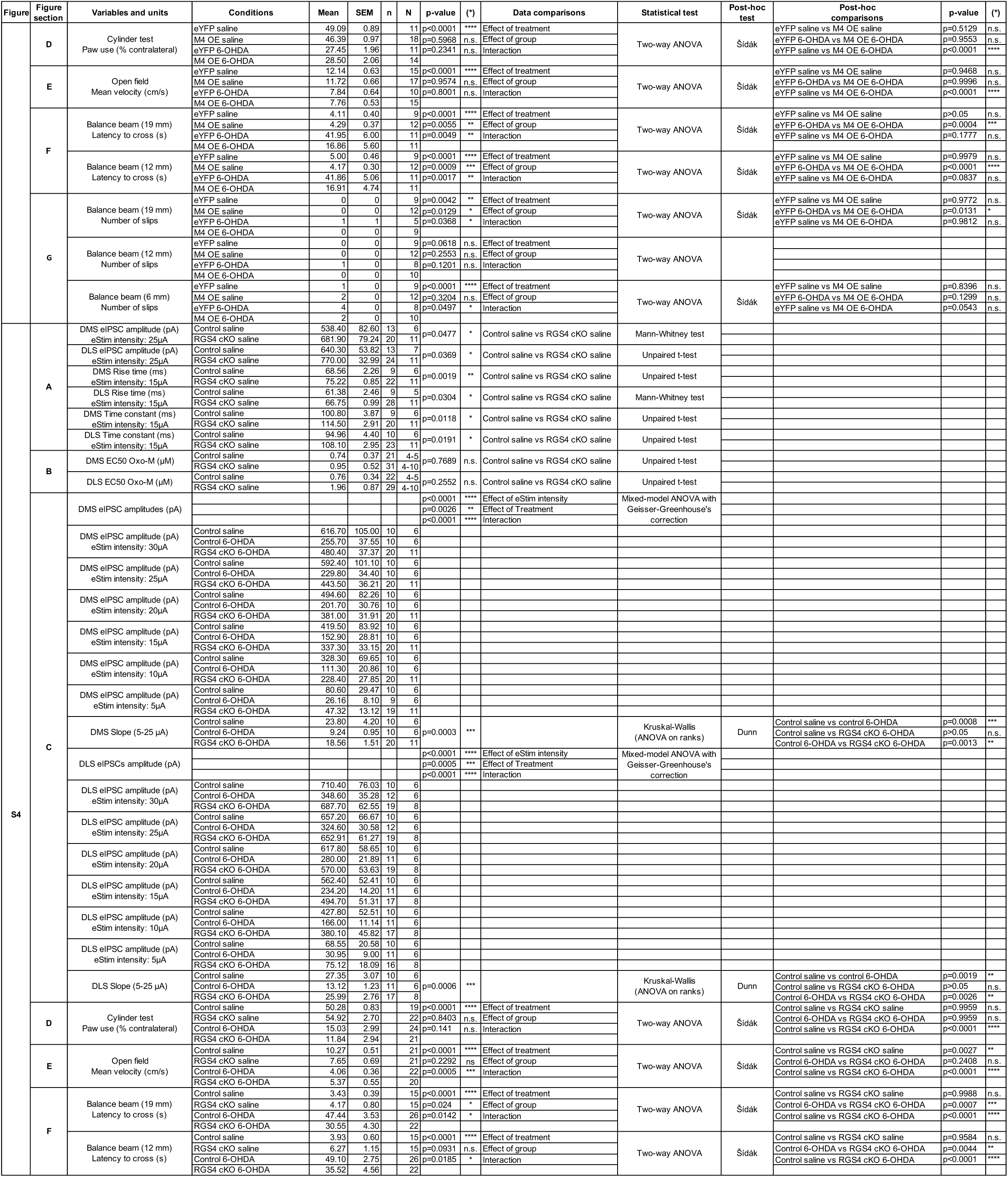

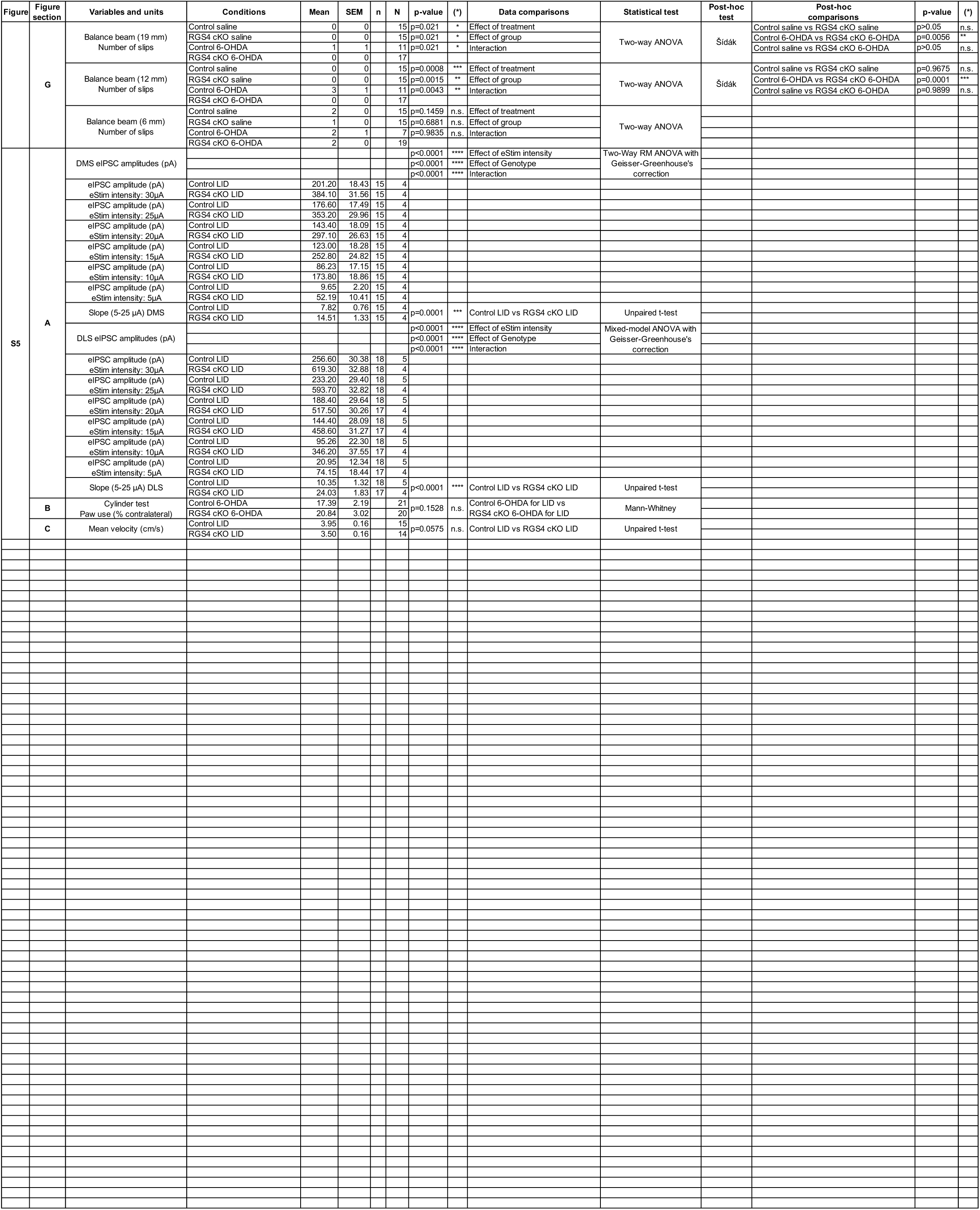

